# Cell deformations generated by dynamic cortical actin waves drive *in vivo* swimming migration

**DOI:** 10.1101/2024.06.11.598427

**Authors:** Cyril Andrieu, Bren Hunyi Lee, Anna Franz

## Abstract

Amoeboid cell migration drives many important developmental and disease-related processes including immune responses and cancer metastasis. Swimming cell migration is a subtype of amoeboid migration observed in cells in suspension *ex vivo.* However, the mechanism underlying swimming migration *in vivo* under physiological conditions is unknown. Using *Drosophila* fat body cells (FBCs) as a model, we show that FBCs actively swim to patrol the pupa. Their stop-and-go random walk is powered through the generation of oscillatory actomyosin waves, rather than persistent actin flows used by cells swimming *in vitro.* These actomyosin waves exert peristaltic compressive forces as they move to the cell rear. This causes cell elongation towards the front to propel the cell forward. In addition, we demonstrate that, unlike in other types of amoeboid migration, all three RhoGTPases, RhoA, Cdc42 and Rac1, are required for FBC migration. They control actin wave formation by regulating actin polymerisation through the formin Dia. Furthermore, RhoA at the cell rear induces actomyosin contractions via Rho kinase and myosin II to generate cell deformations. Importantly, our work reveals that swimming migration is a novel *in vivo* migration mode for rapid and long-range cell dispersal, potentially also used by other cells such as immune cells and cancer cells when encountering an aqueous environment.

## Introduction

Cell motility is key to many biological processes such as embryonic development, immune responses or cancer invasion. Cells can use two main modes of cell migration, mesenchymal migration or amoeboid migration. Mesenchymal migration has been studied extensively for many decades, resulting in a detailed understanding of the mechanisms regulating force generation and force transmission for this migration mode. Mesenchymal migration is generally characterized by a slow migration speed and elongated cell shape with a clear front-rear polarity. A large flat lamellipodium containing branched actin extends the cell at the front. The cell is anchored to the substrate via integrin-dependent focal adhesions. The cell rear then retracts through actomyosin contractions that are accompanied by the dissolution of focal adhesions, thereby propelling the cell body forward (Lauffenburger and Horwitz, 1996, Ridley et al., 2003).

While cells *ex vivo* on flat adhesive substrates usually migrate by mesenchymal migration, cells in more complex 3D environments *ex vivo* and *in vivo* often use alternative migration modes. These have in common that they do not involve lamellipodia and are collectively called amoeboid migration (Paluch et al., 2016). A range of cell types use amoeboid migration including embryonic cells (Ruprecht et al., 2015, Lin et al., 2022, Kardash et al., 2010), immune cells (Liu et al., 2015, Lammermann et al., 2008, O’Neill et al., 2018, Aoun et al., 2020, Garcia-Seyda et al., 2021), and some types of cancer cells (Poincloux et al., 2011, Bergert et al., 2015, Ullo and Logue, 2021). Despite the wide range of amoeboid migration mechanisms employed by these different cell types in various environments, the common features are a rounded cell shape, fast migration speed, low or no cell-substrate adhesion and the dependence on actomyosin-dependent cortical actin flows (Paluch et al., 2016). Whilst lacking lamellipodia, these cells can use other types of protrusions including blebs (Bergert et al., 2015, Blaser et al., 2006) or giant blebs (Ruprecht et al., 2015, Liu et al., 2015, Logue et al., 2015, Ullo and Logue, 2021, Bergert et al., 2015), or completely lack protrusions (Poincloux et al., 2011, O’Neill et al., 2018). Together, these characteristics give plasticity to ameboid migratory cells to adapt and move in different 3D environments, in confined conditions and even in solution.

Sperm, bacteria and many protists can migrate by swimming through liquids using the beating or oscillation of cilia or flagella (Elgeti et al., 2015, Ginger et al., 2008). Moreover, it was recently reported that other cells that lack cilia or flagella, can also swim *in vitro* when placed in solution, including the amoeba *Dictyostelium* (Barry and Bretscher, 2010, Howe et al., 2013, Van Haastert, 2011) and mammalian immune cells such as neutrophils (Barry and Bretscher, 2010, Garcia-Seyda et al., 2021), primary human T lymphocytes (Aoun et al., 2020) as well as RAW 264.7 macrophages (O’Neill et al., 2018). In the case of macrophages and T lymphocytes, force generation during swimming migration is powered by a continuous rearward flow of cortical actin (O’Neill et al., 2018, Aoun et al., 2020). The mechanism underlying the generation of this actin flow differs between these cell types. In swimming T lymphocytes actin polymerization at the cell front is essential to generate the actin flow while actomyosin contractions have a minor role (Aoun et al., 2020). In contrast, in RAW macrophages in liquid, which normally have a low migratory capacity, optogenetic RhoA activation on one side of the side induces swimming migration in the opposite direction. Here, optogenetic RhoA activation induces local actomyosin contractions driving a rearward actin flow (O’Neill et al., 2018). In both cell types, force transmission during swimming is mediated by coupling the cortical actin flow to the rearward treadmilling of transmembrane proteins at the cell surface to exert viscous forces onto the surrounding liquid. This membrane flow is accompanied by anterograde membrane trafficking from the rear to the front of the cell to maintain membrane homeostasis crucial to support the swimming migration (O’Neill et al., 2018, Aoun et al., 2020). In these two studies, specific *in vitro* conditions were used to allow the cells to swim either by using a complex experimental set-up (Aoun et al., 2020) or by inducing the swimming migration with an optogenetic system to generate actomyosin contractions via RhoA activation (O’Neill et al., 2018). These studies have provided important first insights into the mechanisms driving the swimming migration *ex vivo*, especially with regards to force transmission. However, we still have an incomplete understanding of how forces are generated and transmitted to the environment in swimming cells. Moreover, it is still unknown, whether these cells or any other cell types can migrate by swimming in the highly complex 3D environment found *in vivo* and if so, what mechanism they use to swim *in vivo*.

We recently identified the first *in vivo* model of swimming cell migration, in which fat body cells (FBCs) in the *Drosophila* pupa use swimming cell migration to actively move across the hemolymph, the body fluid of the fly, to reach skin wounds (Franz et al., 2018). *Drosophila* FBCs are the equivalent of vertebrate adipocytes which play several crucial roles during insect life including energy storage (Bi et al., 2012, Beller et al., 2010, Gronke et al., 2005, Gronke et al., 2007), humoral immune response (Lemaitre and Hoffmann, 2007, Buchon et al., 2014) and regulation of tumour growth (Kong et al., 2022, Parvy et al., 2019). We found that in pupae, these giant spherical, polyploid cells actively migrate with high directional persistence to reach wounds where they play several local roles in wound healing. We proposed that pupal FBCs use swimming migration involving cortical actin waves and actomyosin contraction (Franz et al., 2018).

*Drosophila* pupal FBCs are, to our best knowledge, the first and currently only *in vivo* model of swimming cell migration. Here, we use this powerful model that allows high-throughput genetic screening and high-resolution live imaging to study the mechanism underlying force generation during *in vivo* swimming migration. We show that, even in absence of a wound, FBCs use swimming migration to patrol the whole pupa. We find that swimming migration is powered by dynamic cortical actomyosin waves at the cell rear resulting in cell deformations which propel the cell forward. We find that actin wave generation and FBC deformations are finely regulated by the small RhoGTPases RhoA, Rac1 and Cdc42 and several of their key downstream regulators to induce actin polymerization and actomyosin contraction during migration.

## Results

### Patrolling fat body cells migrate around the pupa by active cell migration

We have previously shown that FBCs actively migrate towards epithelial wounds with high directional persistence (Franz et al., 2018). To investigate the mechanism underlying the swimming migration of the FBCs in more detail, we first characterised the general migration behaviour of FBCs in the absence of a wound. To follow the migration of FBCs in the whole pupa, we took advantage of the fact that these giant cells have large polyploid nuclei (around 20µm in diameter) to track them in 2D (for ease of data analysis) during 3h-long movies using a widefield microscope. For this, we expressed Nuclear Localisation Signal (NLS)-mCherry exclusively in FBCs by using the UAS-Gal4 system with the FBC-specific Lsp2-Gal4 driver (Deutsch et al., 1989). This new high-throughput automatic nuclear tracking assay allowed us to assess the migration behaviour (including track path, speed, directionality, total displacement) of ∼280-320 cells per pupa in dozens of pupae per condition (in total: thousands of cells per condition; only analysing continuous tracks of 1.5h-3h length). We also used this to measure the mean speed for each pupa from the mean speed of all tracked FBCs. We found that in 16h APF (after pupa formation) old pupa, in the absence of any wound, FBCs were highly migratory (Figure 1A and Movie 1A). In the head and thorax, FBCs migrated at a high speed (mean speed: 3.2 and 3µm/min, respectively; Figure 1A and C). This fast migration led to the displacement of FBCs inside the pupa over very long distances of up to 153 micrometres in 3h (Suppl. Figure 1A). Interestingly, in the abdomen, FBCs migrated with a significantly lower speed (mean speed: 2.3µm/min; Figure 1A and C and Movie 1A) and lower straightness (Suppl. Figure 1B) than in the head and thorax. In the whole pupa, the mean speed for individual FBC tracks was highly variable, ranging from 1µm/min to 5.3µm/min (Suppl. Figure 1C). Moreover, in the head and thorax, individual FBC tracks frequently alternated between periods of slower and faster migration (Suppl. Figure 1D-F; mean maximum speed: 6-7µm/min, highest maximum speed: 11-13µm/min, mean minimum speed: 0.4-0.7µm/min, lowest minimum speed: ∼0.02-0.06µm/min). This resembles a stop-and-go or run-and-tumble mode of migration resulting in random walk (Zhang et al., 2023, Miller et al., 2002).

**Figure 1.**
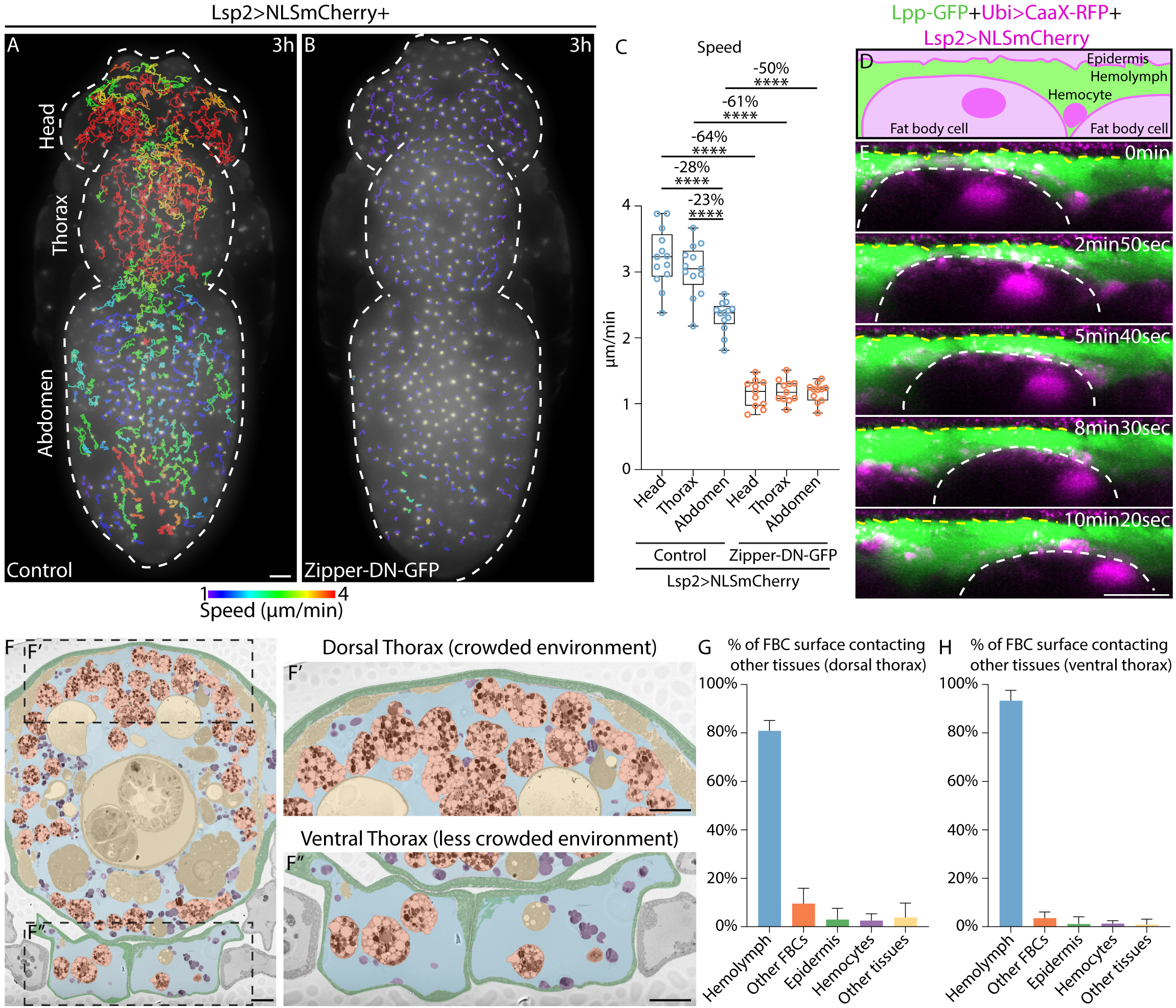
*Drosophila* fat body cells patrol the pupa by swimming migration. (A-B) Widefield timelapse images of the dorsal view of *Drosophila* pupae expressing Lsp2-Gal4+UAS-NLS-mCherry+control (A) or +UAS-Zipper-DN-GFP (B). Only continuous 1h30-3h long migration tracks are shown color-coded according to their mean speed. See also Movie 1. (C) Quantification of mean FBC speed from (A, B) in the head, thorax and abdomen for control (n:13pupae) and Zipper-DN-GFP (n:11pupae). Showing mean speed for each pupa calculated from the mean speed of each of its 1h30-3h-long tracks. Ordinary one-way ANOVA test with multiple comparisons, ****P<0.0001. (D-E) Schematic (D) and XZ view of confocal time lapse images (E) of two FBCs swimming in hemolymph near the epidermis in a Lpp-GFP+Ubi>CaaX-RFP+Lsp2-Gal4+UAS-NLS-mCherry pupa (hemolymph in green; epidermis in magenta above yellow-dashed line on the top; FBCs in faint magenta with bright magenta nuclei, FBC outlined with white dotted line; note presence of a hemocyte containing a bright magenta object). See also Movie 2. (F-F’’) Methylene-blue stained, transverse, semi-thick section of the pupal thorax with false-colored tissues showing FBCs (orange), hemolymph (blue), epidermis (green), hemocytes (purple) and other tissues (yellow). Magnified images of dorsal thorax (F’) and ventral thorax (F’’). (G-H) Quantification of percentage of FBC surface contacting other tissues in dorsal (G) and ventral (H) thorax (n:7cells each) from (F’-F”). Using mean for each cell calculated from 15-20 serial sections. Scale bars, 100 µm (A-B), 20 µm (E), 50 µm (F-F’’)

Next, we wanted to determine whether this motility is indeed due to active migration rather than a consequence of FBCs being moved around passively, e.g. by potential hemolymph flows that might be generated through muscle or heart contractions. To test this, we expressed a dominant-negative (DN), GFP-tagged version of Zipper specifically in FBCs. Zipper is the heavy chain of myosin II, known to be essential for actomyosin contractions and cell migration in other cell types (Majumder et al., 2012, Yolland et al., 2019). We found that expression of Zipper-DN-GFP led to a strong, significant reduction in migration speed (50-64% lower) in all three compartments (Figure 1B-C and Movie 1B) compared to the control (Figure 1A and C and Movie 1A). This suggests that the observed movement of FBCs in all areas of the pupal body is mostly driven by active cell migration.

### Fat body cells migrate in the hemolymph by swimming migration

In our previous study, we suggested that FBCs use directed swimming migration to go to wounds in the ventral thorax (Franz et al., 2018). This part of the pupa only contains small numbers of FBCs that are sparsely distributed in the hemolymph (Figure 1F and F’’) which is suboptimal for large scale analysis of FBC migration. Hence, we decided to use the dorsal head and thorax, where there are many fast migrating FBCs at higher density (Figure 1F-F’), to study the mechanism of FBC migration further. To see whether, in these more crowded regions, FBCs also use contact-independent swimming migration, we first visualised the migration of FBCs in relation to the epidermis (labelled with ubiquitously expressed CaaX-RFP) and the hemolymph (labelled with Lpp-GFP). We observed that in most cases, FBCs (labelled strongly with NLS-mCherry and weakly with CaaX-RFP) did not make any close contact with the epidermis during migration (Figure 1D-E and Movie 2). The cells always kept their spherical shape rather than flattening, and moved within the hemolymph, confirming that FBCs swim independent of cell-epidermis contacts. Moreover, we quantified in methylene blue-stained, serial sections of the pupal thorax, how much of the FBC surface is in contact with hemolymph versus various tissues. In the dorsal thorax, 80% of the surface of FBCs was surrounded by hemolymph with only minor contact to other FBCs, the epidermis, hemocytes (*Drosophila* macrophages) or other pupal tissues (Figure 1F’ and G). It is possible that these transient, minor cell-cell contacts might contribute to FBC migration in these crowded regions. However, we have previously shown that in the much less crowded ventral thorax where many FBCs are exclusively surrounded by hemolymph (Figure 1F’’ and H and (Franz et al., 2018)), FBCs migrate at very similar speed (mean speed: ∼3.5µm/min (Franz et al., 2018)) to the one of FBCs in the head and thorax (Figure 1C). Together, our results indicate that, similar to in the less crowded ventral thorax, the migration of the FBCs in the more crowded environment of the dorsal head and thorax is by swimming migration with limited cell-cell contact.

### Contractile cortical actin waves drive cell deformations in the cell rear of migrating FBCs

In amoeboid migration and *in vitro* swimming migration, rearward cortical actin flows usually generate the main force to power migration (Ruprecht et al., 2015, Liu et al., 2015, Bergert et al., 2015, Aoun et al., 2020, Lin et al., 2022). We previously described that FBCs migrating towards wounds in the ventral thorax generate actin waves in their cortex (Franz et al., 2018). We next tested if in the absence of a wound similar actin waves are also associated with the general migration of FBCs in the dorsal head and thorax. To visualize actin dynamics, we performed *in vivo* live imaging using various actin markers. Several previous studies, including one in *Drosophila* nurse cells in egg chambers (Spracklen et al., 2014), showed that the expression of some actin markers can negatively affect the actin network. Similarly, we found that expression of GMA-GFP (Edwards et al., 1997) and Utrophin-GFP (Burkel et al., 2007) impacted FBC migration, with a significant increase or decrease in FBC speed, respectively (Suppl. Figure 2A and C-E). However, speed was not significantly altered by expression of LifeAct-GFP (Riedl et al., 2008) (Suppl. Figure 2A-B and E).

Based on these results, we decided to use LifeAct-GFP to study actin dynamics in swimming FBCs. We found that periodic actin waves, that were restricted to the FBC cortex, were present in all FBCs in the pupal head (Figure 2A and Movie 3, red arrowheads pointing at visually detectable actin waves defined by a local increase followed by a decrease in intensity as seen at 5min, 7min, 9min30, 12min30, 14min and 16min30). The waves usually moved towards the rear of the cell where they induced clear local deformations of the cell surface followed by an elongation of the cell body (measured as an increase in the cell aspect ratio) in the opposite direction (Figure 2A-C and Movie 3). Peaks of actin waves were often followed within ∼1.5min by peaks in aspect ratio of FBCs (Figure 2C, red arrowheads and blue arrowheads, respectively) suggesting that actin waves induce cell elongations. Strikingly, as actin waves moved along the cell surface towards the rear, they often locally compressed the cell surface in a belt-like manner. This resulted in the cell rear retracting and in the fast propulsion of the cell body in the opposite direction (Figure 2D, D’, E, red arrowheads showing actin wave and blue arrows pointing at concave cell surface deformations, and Movie 4).

**Figure 2.**
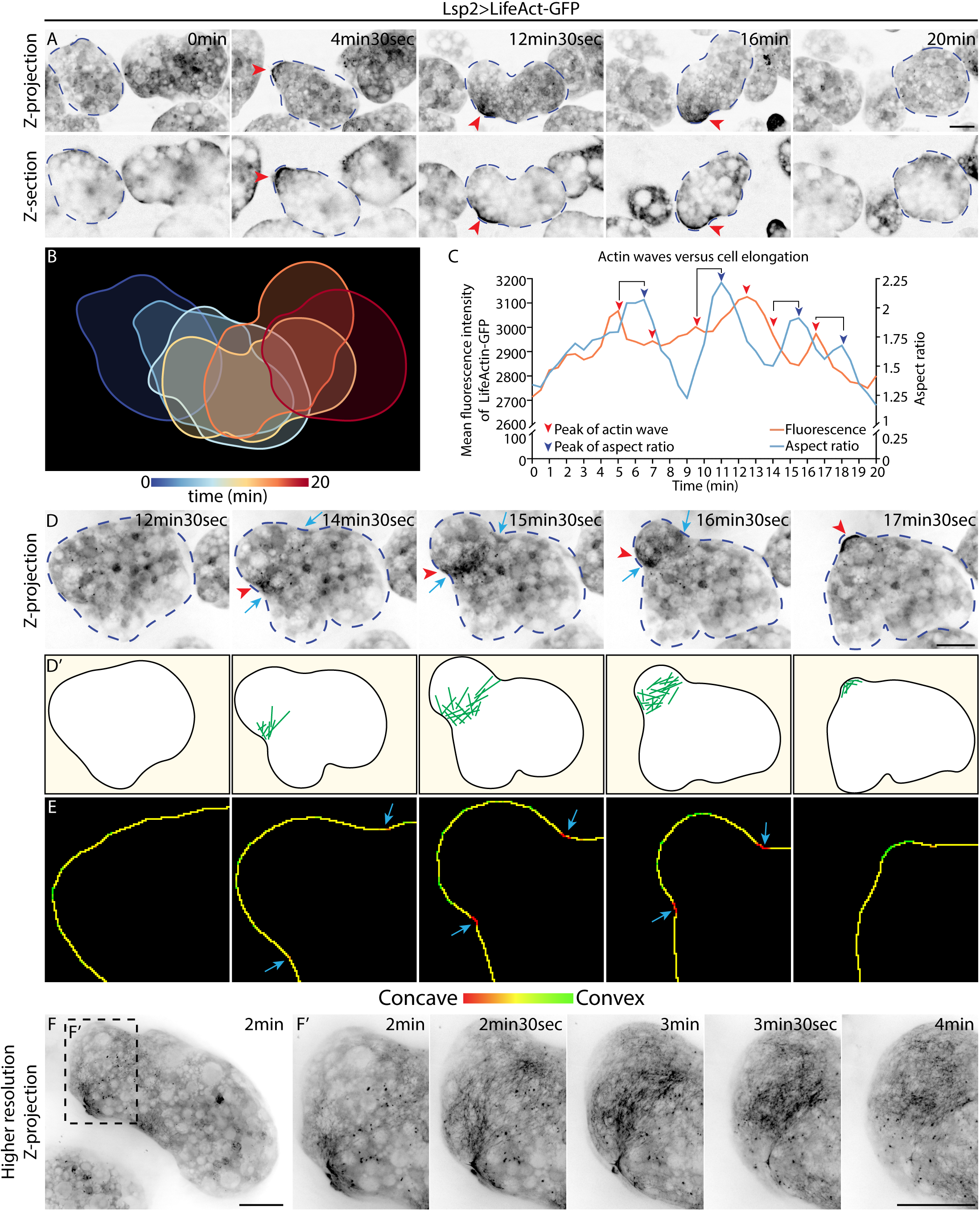
Cortical actin waves lead to cell deformations at cell rear to propel fat body cells during migration. (A) Confocal time-lapse images of actin dynamics in migrating FBCs expressing Lsp2-Gal4+UAS-LifeAct-GFP in Z-projection (top images) and in Z-section deeper in the FBCs (bottom images). Red arrowheads pointing at consecutive actin waves in the back of a migrating FBC (outlined with blue dotted line). See also Movie 3. (B) Color-coded time projection of cell outline from 6 time points from time-lapse movie from (A) to visualise deformations of the FBC during swimming migration. (C) Quantification of the mean fluorescence intensity of LifeAct-GFP and the cell aspect ratio of the migrating FBC from (A). Red arrowheads point at the peaks of the actin waves assessed visually in Movie 3 which also match the peaks in mean fluorescence intensity (with the exception of the 4^th^ and 5^th^ wave that are only apparent as separate actin waves when assessed visually) and blue arrowheads point at peaks in cell aspect ratio. (D, D’) Confocal time-lapse images (D) and schematic (D’) of a Lsp2+UAS-LifeAct-GFP expressing FBC. Red arrowheads show actin wave moving to the rear and blue arrows point at FBC cell surface compressions (D). Actin shown in green in schematic (D’). See also Movie 4. (E) Magnified images from (D) showing cell outline with concave (red) and convex (blue) surface areas at different time points. Green arrows point at concave FBC deformations. (F-F’) ‘Higher resolution’ time-lapse images of a Lsp2+UAS-LifeAct-GFP expressing FBC. Magnified view of F in (F’). See also Movie 5. Scale bars, 20 µm.

To look at the actin network in the waves in more detail, we performed high resolution *in vivo* live imaging using a SoRa spinning disc microscope with a 40x silicone lens with a magnification of 2.8 (hereafter referred to as ’higher resolution imaging’), which resulted in increased signal to noise ratio with lower background fluorescence and increased the definition of the actin network. We observed that areas in the wave periphery seemed to contain more individual actin filaments, while the wave centre contained a denser actin meshwork (Figure 2F-F’ and Movie 5, observed in 8/8 cells). Actin waves moved along the cell surface to nearby areas previously devoid of actin filaments. Overall, the actin meshwork seemed highly dynamic and usually did not cover the entire cortex of the cell, with large parts of the surface being seemingly devoid of actin (Figure 2F-F’ and Movie 5, observed in 8/8 cells). In contrast, expression of the actin marker Utrophin-GFP, which, as we mentioned before reduced migration speed (Suppl. Figure 2A, D and E), resulted in an unusually stable, dense actin meshwork. It covered most of the cortex, contained many thick actin bundles and some actin was also found in circular structures deeper inside the cell (Suppl. Figure 2H) which were not observed with LifeAct-GFP and GMA-GFP (Suppl. Figure 2F-G). Together, this suggests that the FBC actin network has to be dynamic to allow efficient FBC migration.

Mesenchymal and amoeboid migration often involves the formation of protrusions in the cell front such as lamellipodia and blebs, respectively. In contrast, when assessing actin dynamics and cell shapes during FBC migration, we never observed any such actin-rich protrusions or blebs in the cell front which always kept its rounded shape (Movies 3-5).

Altogether, these data show that FBCs migrate within the aqueous environment of the hemolymph using amoeboid swimming. They maintain a rounded cell shape without forming protrusions in the cell front. Instead, they use contractile cortical actin waves to generate forces to deform the cell at the rear and to propel the cell body forward and swim.

### The small RhoGTPase Rho1 is crucial for FBC migration by regulating actin wave formation and cell deformations

Next, we wanted to dissect the molecular mechanism regulating force generation during FBC migration. For this, we took advantage of our newly developed automatic nuclear tracking assay that allows assessing the migration speed of thousands of cells from dozens of pupae per genotype in high-throughput (Figure 1A-C).

We started with the small RhoGTPase RhoA which plays a key role during amoeboid migration by controlling the formation of a retrograde actin flow and inducing contractility at the cell rear (Kardash et al., 2010, Lin et al., 2022, Poincloux et al., 2011). Moreover, photoactivation of RhoA in RAW macrophages is sufficient to induce *in vitro* swimming migration (O’Neill et al., 2018). To determine whether RhoA (Rho1 in *Drosophila*) is involved in FBC migration, we expressed a dominant negative form of Rho1 (Rho1-N19) specifically in FBCs. This resulted in a strong, significant 53% reduction in FBC migration speed (Figure 3A-B, D and Movie 6A-B). To confirm this result, we also expressed Rho1 RNAi specifically in FBCs which also significantly reduced FBC migration speed (Figure 3A, C-D and Movie 6A, C). This suggests that Rho1 plays a key role in the swimming migration of FBCs.

**Figure 3.**
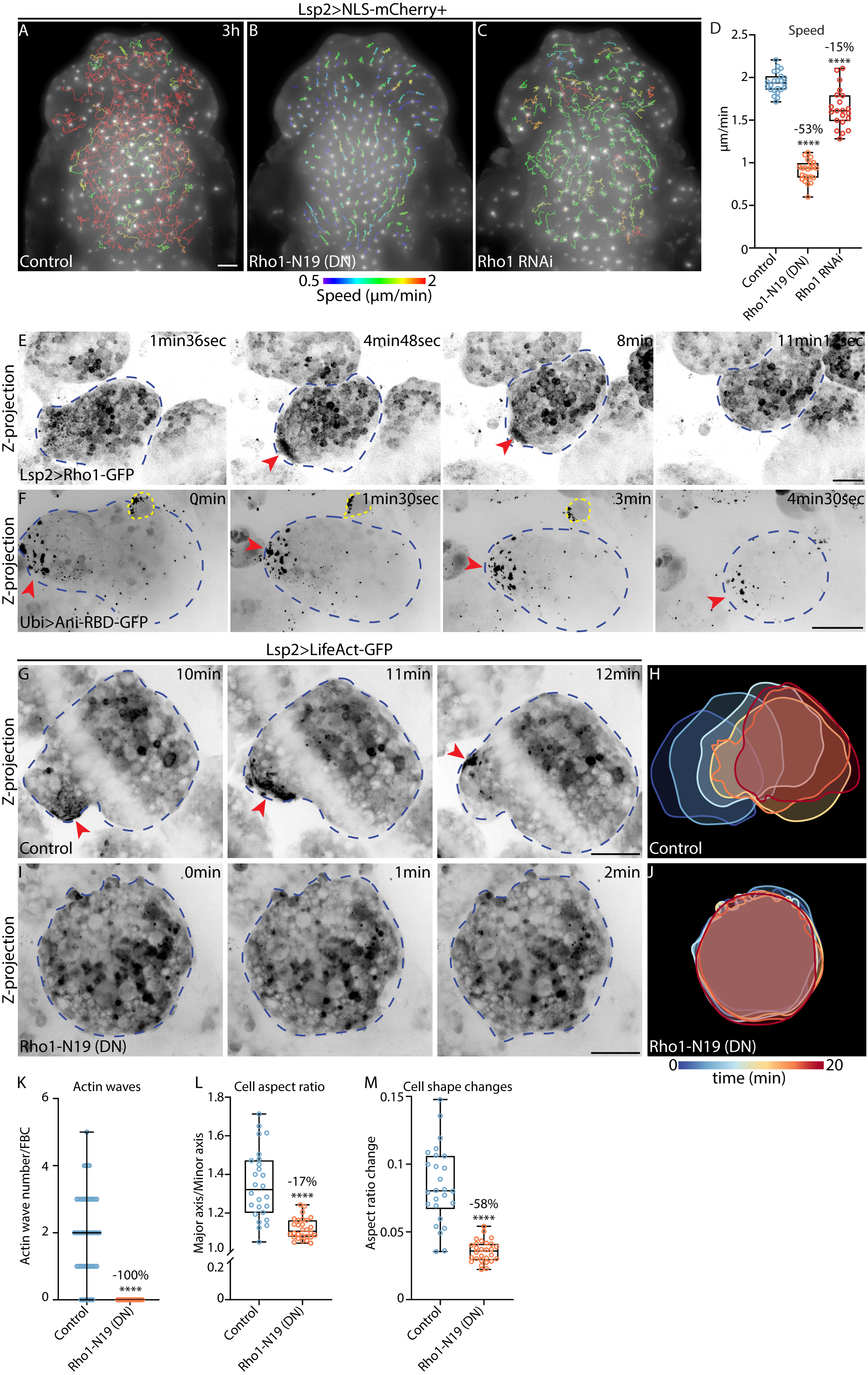
Rho1 is needed for actin wave formation and cell deformations during fat body cell swimming migration. (A-C) Widefield timelapse images of the dorsal head and thorax of *Drosophila* pupae expressing Lsp2-Gal4+UAS-NLS-mCherry+control (A), +UAS-Rho1-N19 (dominant negative form of Rho1) (B) or +UAS-Rho1 RNAi (C). 1h30-3h-long migration tracks are shown color-coded according to their mean speed. See also Movie 6. (D) Quantification of mean FBC speed from (A-C). Control (n:23pupae), Rho1-N19 (n:23pupae) and Rho1 RNAi (n:21pupae). Showing mean speed for each pupa calculated from the mean speed of each of its 1h30-3h-long tracks. Ordinary one-way ANOVA test with multiple comparisons **** p<0.0001. (E) Confocal time-lapse images of a swimming FBC expressing Lsp2-Gal4+UAS-Rho1-GFP. Red arrowhead points at the accumulation of Rho1-GFP in the rear of a migratory FBC. Blue dotted line outlines FBC. See also Movie 7A. (F) Confocal time-lapse images of a swimming FBC in a pupa expressing Ani-RBD-GFP under the ubiquitin p63E promoter. Red arrowhead points at the accumulation of Ani-RBD-GFP seen as punctae in the rear of a migratory FBC. Blue dotted line outlines FBC and yellow dotted line outlines a hemocyte. See also Movie 7B. (G-J) Confocal time-lapse images of actin dynamics (G and I) and color-coded time projection of cell outlines from several time points (H and J) from FBCs expressing Lsp2-Gal4+UAS-LifeAct-GFP+control (G, H) or +UAS-Rho1-N19 (I, J). Red arrowhead points at actin wave and blue dotted line outlines FBC. Note that white line in image G is the shadow formed by a cuticle fold. See also Movie 8. (K) Quantification of number of actin waves seen during 20min-long movies from (G and I) in FBCs expressing Lsp2-Gal4+UAS-LifeAct-GFP+control (n:59cells) and +UAS-Rho1-N19 (n:52cells). Mann-Whitney test ****p<0.0001. (L, M) Quantification of mean aspect ratio (L) and mean aspect ratio change over time (M) from (G and I) of FBCs expressing Lsp2-Gal4+UAS-LifeAct-GFP+control (n:26cells) and +UAS-Rho1-N19 (n:52cells). Unpaired t test ****p<0.0001. Scale bars, 100 µm (A-C), 20 µm (E-G and I).

We next studied the subcellular localization of Rho1 by expressing Rho1-GFP in the FBCs with the FBC-specific Lsp2-Gal4 driver. We observed an accumulation of Rho1-GFP in the cell rear of migrating FBCs (Figure 3E, red arrowhead, and Movie 7A) where actin waves and cell deformation are usually found (Figure 2A and D). We also used the Rho1 GTP binding domain of Anillin fused to GFP (Ani-RBD-GFP) previously used to assess the localization of the active form of Rho1 (Munjal et al., 2015). Since Ani-RBD-GFP expression is under the control of the ubiquitin p63E promoter, it is expressed in many pupal tissues. Similar to Rho1-GFP, Ani-RBD-GFP also accumulated in the cell rear of migrating FBCs (Figure 3F, red arrowhead and Movie 7B). This suggests that during FBC migration Rho1 accumulates and gets activated at the cell rear where it might control actin waves and cell deformations.

To study how Rho1 regulates FBC migration further, we next used LifeAct-GFP expression to test whether the block of Rho1 function affects actin wave formation and cell deformations. In control FBCs, between 0 and 5 actin waves with an average of two actin waves formed in 20 minutes (Figure 3G, red arrowhead, K and Movie 8A). In contrast, Rho1-N19 expression completely abolished actin wave formation (Figure 3I, K and Movie 8B). Moreover, while control FBCs were more deformed with undulating circular cell shapes and displayed dynamic cell deformations (Figure 3H, L and Movie 8A), Rho1-N19-expressing FBCs were more circular in shape throughout the length of the movies and had a significant lower aspect ratio (Figure 3J, L and Movie 8B). To quantify these dynamic changes in cell shape normally seen in the control, we calculated the change in aspect ratio every minute and we found a strong, significant reduction of mean aspect ratio change upon Rho1-N19 expression (Figure 3M). Thus, Rho1 is a key regulator of FBC swimming migration by controlling the formation of actin waves and regulating dynamic cell shape changes.

### Actomyosin contractility drives FBC deformations during swimming migration

Having found that Rho1 regulates actin wave formation and FBC deformations, we next wanted to identify the downstream effectors involved in this regulation. RhoA is generally known to regulate actomyosin contractility through Rho kinase (Rok) and myosin II (Ridley, 2001, Ridley, 2015). Actomyosin contractility, in turn, drives the generation of a retrograde actin flow in amoeboid migration by myosin pulling on the cortical actin at the cell rear (Shih and Yamada, 2010, Poincloux et al., 2011). We hypothesized that in migrating FBCs, Rho1 might similarly regulate actomyosin contractility through Rok and myosin II to generate contractile actin waves and FBC deformations. Hence, we blocked Rok, with two RNAi lines, and myosin II, with a dominant-negative, phosphorylation-insensitive form (sqh-AA) of the light chain of myosin II (Sqh), a Sqh RNAi line, and with the dominant negative form of Zipper (Zipper-DN-GFP), the heavy chain of myosin II. In all conditions we observed a strong, significant reduction in FBC migration speed (Figure 4A-G and Movie 9). This suggests that Rok and myosin II play an important function in FBC migration and points further at the importance of Rho1 in FBC migration.

**Figure 4.**
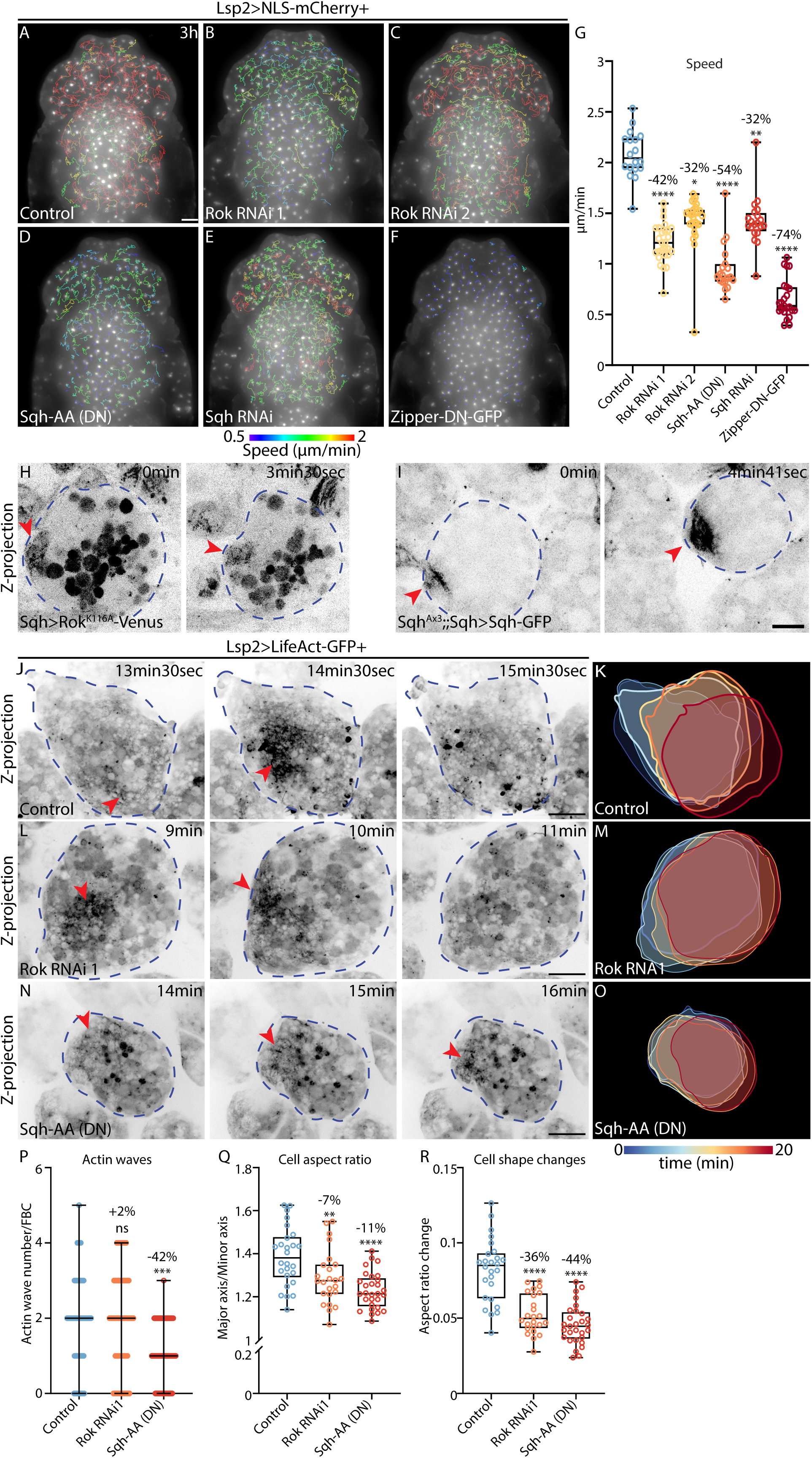
Actomyosin contractility is essential for fat body cell deformations during swimming migration. (A-F) Widefield timelapse images of the dorsal head and thorax of *Drosophila* pupae expressing Lsp2-Gal4+UAS-NLS-mCherry+control (A), +UAS-Rok RNAi1 (B), +UAS-Rok RNAi2 (C), +UAS-Sqh-AA (DN) (D), +UAS-Sqh RNAi (E) or +UAS-Zipper-DN-GFP (F). 1h30-3h-long migration tracks are shown color-coded according to their mean speed. See also Movie 9. (G) Quantification of mean FBC speed from (A-F). Control (n:18pupae), Rok RNAi (n:27pupae), Rok RNAi2 (n:27pupae), Sqh-AA (DN) (n:19pupae), Sqh RNAi (n:20pupae) and Zipper-DN-GFP (n:23pupae). Showing mean speed for each pupa calculated from the mean speed of each of its 1h30-3h-long tracks. Kruskal-Wallis test with multiple comparisons *p=0.0128, **p=0.0061 and ****p<0.0001. (H) Confocal time-lapse images of migratory FBC expressing Rok^K116A^-Venus under the sqh promoter. Blue dotted line outlines FBC and red arrowhead points at the accumulation of Rok^K116A^-Venus in the FBC rear. See also Movie 10A. (I) Confocal time-lapse images of migratory FBC expressing Sqh-GFP under the sqh promoter in a sqh mutant background. Blue dotted line outlines FBC and red arrowhead points at the accumulation of Sqh-GFP in the FBC rear. See also Movie 10B. (J-O) Confocal time-lapse images of actin dynamics (J, L and N) and color-coded time projection of cell outlines from several time points (K, M and O) from FBCs expressing Lsp2-Gal4+UAS-LifeAct-GFP+control (J, K), +UAS-Rok RNAi1 (L, M) or +UAS-Sqh-AA (N, O). Red arrowhead points at actin wave and blue dotted line outlines FBC. See also Movie 12. (P) Quantification of number of actin waves seen during 20min-long movies from (J, L and N) in FBCs expressing Lsp2-Gal4+UAS-LifeAct-GFP+control (n:55cells), +UAS-Rok RNAi1 (n:62cells) and +UAS-Sqh-AA (DN) (n:64cells). Kruskal-Wallis test with multiple comparisons, ns p>0.9999 and *** p:0.0006. (Q, R) Quantification of mean aspect ratio (Q) and mean aspect ratio change over time (R) from (J, L and N) of FBCs expressing Lsp2-Gal4+UAS-LifeAct-GFP+control (n:26cells) or +UAS-Rok RNAi1 (n:24cells) or +UAS-Sqh-AA (DN) (n:29). Ordinary one-way ANOVA test with multiple comparisons, ** p:0.0075 and **** p<0.0001. Scale bars, 100 µm (A-F), 20 µm (H-J, L and N).

Next, we studied the subcellular localization of Rok and myosin II during FBC migration. We used lines in which expression of Rok^K116A^-Venus or Sqh-GFP is driven in several cell types by the sqh promoter. Rok^K116A^ is a mutant variant with a disrupted catalytic activity to avoid high-levels of Rok activity in the cell. The Sqh-GFP line is in a sqh mutant background to avoid potential negative side effects caused by Sqh overexpression. Strikingly, Rok^K116A^-Venus and Sqh-GFP accumulated in the cortex at the rear of migrating FBCs (Figure 4H-I, red arrowhead and Movie 10), where we usually also find actin waves (Figure 2A). Moreover, higher resolution imaging showed that Sqh-GFP formed a meshwork closely associated with the actin wave (Movie 11), suggesting that myosin II might pull on the actin to concentrate the actin to generate strong actin waves.

To test this hypothesis, we examined actin wave formation in FBCs when we blocked Rok and Sqh function. Interestingly, in both cases FBCs still had actin waves although they appeared less strong than in the control (Figure 4J, L, N, P and Movie 12). Moreover, in contrast to Rok RNAi, the expression of Sqh-AA significantly decreased the number of actin waves by 50% (Figure 4P). This difference between Rok-RNAi and Sqh-AA could be explained either by myosin II getting activated by Rok as well as another kinase, or by the loss of function effect of Sqh-AA being stronger than the one of Rok-RNAi. Both hypotheses are also supported by the stronger reduction in migration speed that we found upon Sqh-AA expression compared to Rok RNAi (Figure 4G). Together, these data suggest that actomyosin contraction, regulated by Rok and myosin II, at least in part, contributes to the regulation of actin waves.

Next, we hypothesized that myosin-driven contraction of the cortical actin during actin wave formation would be particularly important for inducing cell deformations by locally pulling the cell surface inwards. Indeed, we found that the loss of function of Rok and myosin II affected the cell shape of migrating FBCs with a significant reduction in aspect ratio of FBCs (Figure 4Q). Moreover, there was a strong decrease in cell shape changes over time (Figure 4K, M, O, R and Movie 12).

In summary, actomyosin contractions in the cell rear regulated by Rok and myosin II are crucial for FBC deformations and contribute, to a lesser extent, to actin wave regulation during swimming migration.

### The formin Dia regulates actin wave formation by driving actin polymerization

In mesenchymal and amoeboid cell migration, in addition to regulating actomyosin contraction in the cell rear, RhoA is known to induce actin polymerization via the formin Dia (Ridley, 2001, Ridley, 2015). To test whether Dia-dependent actin polymerization is important for actin wave formation and cell migration in FBCs, we affected Dia function through FBC-specific expression of two different RNAi lines as well as of a constitutive active form of Dia (Dia-ΔDAD-GFP). All of these resulted in a significant, moderate to strong reduction in FBC migration speed (Figure 5A-E and Movie 13), suggesting that Dia is required for FBC migration. Moreover, Dia RNAi strongly reduced actin wave numbers (Figure 5F, H, N and Movie 14) and resulted in less elongated cell shapes and strongly reduced cell shape changes (Figure 5G, I, O and P). In contrast, Dia overactivation resulted in a strongly increased cortical actin meshwork covering most of the cortex and containing many swirls of parallel actin bundles (Figure 5J-L, red arrowheads and Movie 15). These highly dynamic actin swirls moved along the whole FBC cortex and appeared different to actin waves in the control (Movie 15). Dia overactivation also induced the formation of some type of cell surface protrusions containing strongly concentrated actin (Figure 5L, blue arrows). The overall cell shape was also impacted upon Dia overactivation, with FBCs being more circular throughout time and deforming less than in the control (Figure 5K, M, Q and R).

**Figure 5.**
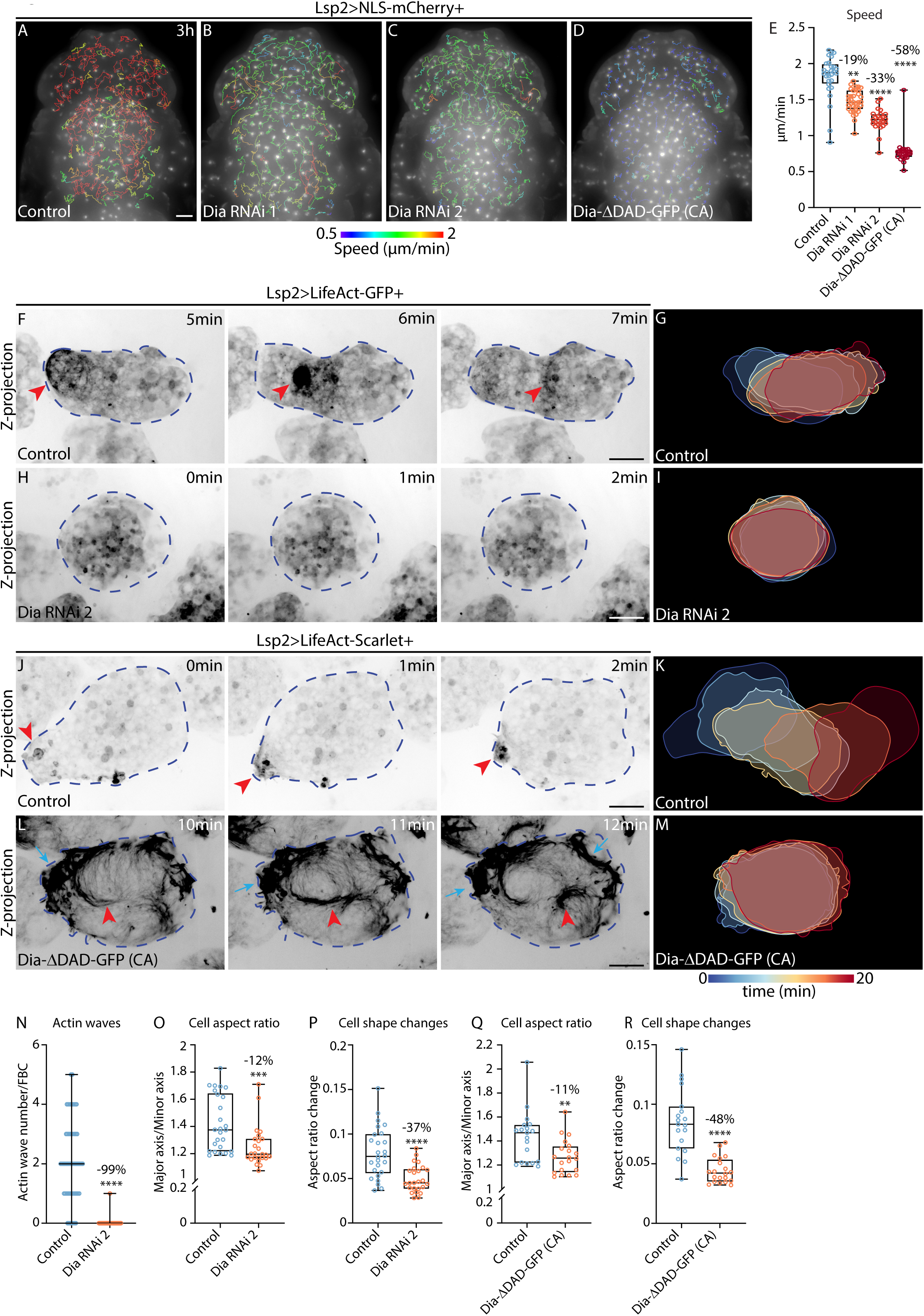
Actin waves are regulated by the formin Dia. (A-D) Widefield timelapse images of the dorsal head and thorax of *Drosophila* pupae expressing Lsp2-Gal4+UAS-NLS-mCherry+control (A), +UAS-Dia RNAi1 (B), +UAS-Dia RNAi2 (C) or +UAS-Dia-ΔDAD-GFP (CA) (D). 1h30-3h-long migration tracks are shown color-coded according to their mean speed. See also Movie 13. (E) Quantification of mean FBC speed from (A-D). Control (n:28pupae), Dia RNAi1 (n:35pupae), Dia RNAi2 (n:25pupae) and Dia-ΔDAD-GFP (CA) (n:26pupae). Showing mean speed for each pupa calculated from the mean speed of each of its 1h30-3h-long tracks. Kruskal-Wallis test with multiple comparisons **p=0.0065 and ****p<0.0001. (F-M) Confocal time-lapse images of actin dynamics (F, H, J, L) and color-coded time projection of cell outlines from several time points (G, I, K, M) from FBCs expressing Lsp2-Gal4+UAS-LifeAct-GFP+control (F, G) or +UAS-Dia RNAi2 (H, I), or expressing Lsp2-Gal4+UAS-LifeAct-Scarlet+control (J, K) or +UAS-Dia-ΔDAD-GFP (L, M). Red arrowheads point at actin waves (F and J) or at dynamic actin swirls (L), blue arrows point at cell surface protrusions containing strongly concentrated actin (L) and blue doted lines outline FBCs. See also Movies 14 (F, H) and 15 (J, L). (N) Quantification of number of actin waves seen during 20min-long movies from (F, H) in FBCs expressing Lsp2-Gal4+UAS-LifeAct-GFP+control (n:55cells) or +UAS-Dia RNAi2 (n:59cells). Mann-Whitney test, **** p<0.0001. (O-R) Quantification of mean aspect ratio (O, Q) and mean aspect ratio change over time (P, R) from (F, H, J and L) of FBCs expressing Lsp2-Gal4+UAS-LifeAct-GFP+control or +UAS-Dia RNAi2 (O, P) or expressing Lsp2-Gal4+UAS-LifeAct-Scarlet+control or +UAS-Dia-ΔDAD-GFP (Q, R). Control (n:26cells) and Dia RNAi2 (n:25cells) (O, P). Control (n:19cells) and Dia-ΔDAD-GFP (n:20cells) (Q, R). Mann-Whitney test, ***p=0.0002 (O, Q). Unpaired t test, **** p<0.0001 (P, R). Scale bars, 100 µm (A-D), 20 µm (F-H and J-L).

These data suggest that actin polymerization regulated by Dia is crucial for actin wave formation in migrating FBCs. Moreover, they are consistent with our other data showing that, similar to Dia overactivation, Utrophin-GFP expression stabilizes the actin network and reduces migration speed (Suppl. Figure 2A, D-F and H). This suggests that the cortical actin mesh has to be dynamic and tightly controlled to allow normal FBC migration. Finally, our data confirm that actin wave formation is crucial for the generation of FBC deformations and FBC migration.

### FBC swimming migration is also dependent on Cdc42 and Rac1 activity

In classic mesenchymal cell migration, RhoA regulates actomyosin contraction in the cell rear, while the other two small RhoGTPases, Rac1 and Cdc42, are thought to regulate the formation of lamellipodia and filopodia in the cell front via branched actin formation, respectively (Ridley, 2001, Ridley, 2015). Since amoeboid migration usually does not involve lamellipodia or filopodia, it is generally believed that Rac1, Cdc42 and branched actin formation are mostly dispensable. Indeed, the amoeboid migration of *Drosophila* germ cells (Kunwar et al., 2003, Lin et al., 2020) and *in vitro* swimming migration of the RAW macrophages upon optogenetic RhoA activation (O’Neill et al., 2018) do not rely on Rac1 function. However, other examples of amoeboid migration driven by blebbing in the cell front, including zebrafish germ cells, rely on actin polymerization controlled by Rac1 (Kardash et al., 2010). Hence, we decided to test if Rac1 and Cdc42 might be involved in FBC migration. Interestingly, we found that FBC-specific expression of dominant-negative forms of Cdc42 (Cdc42-N17) and Rac1 (Rac1-N17), and expression of Cdc42 RNAi and Rac1 RNAi strongly decreased migration speed of FBCs in the head and thorax (Figure 6A-H and Movie 16). This suggests, that in addition to Rho1, Cdc42 and Rac1 are also key regulators of the swimming migration of FBCs.

**Figure 6.**
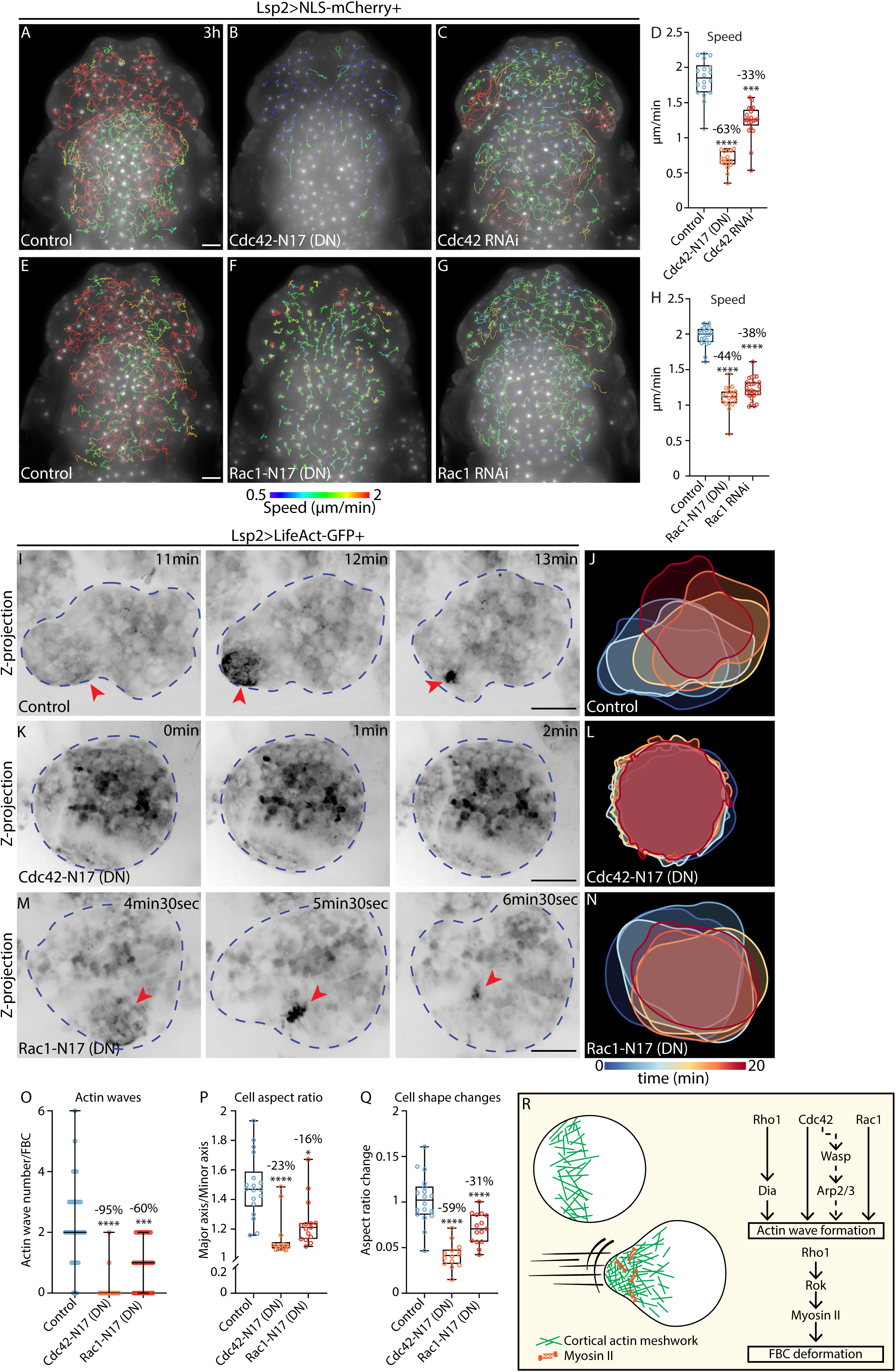
Cdc42 and Rac1 control fat body cell migration through the regulation of actin waves and cell deformations. (A-C, E-G) Widefield timelapse images of the dorsal head and thorax of *Drosophila* pupae expressing Lsp2-Gal4+UAS-NLS-mCherry+control (A), +UAS-Cdc42-N17 (DN) (B) or +UAS-Cdc42 RNAi (C) and +control (E), or +UAS-Rac1-N17 (DN) (F) or UAS-Rac1 RNAi (G). 1h30-3h-long migration tracks are shown color-coded according to their mean speed. See also Movie 16. (D, H) Quantification of mean FBC speed from (A-C and E-G). Control (n:18pupae), Cdc42-N17 (n:17pupae) and Cdc42 RNAi (n:21pupae) (D) and control (n:22pupae), Rac1-N17 (n:19pupae) and Rac1 RNAi (n:26pupae) (H). Showing mean speed for each pupa calculated from the mean speed of each of its 1h30-3h-long tracks. Kruskal-Wallis test with multiple comparisons, ***p=0.0008 and ****p<0.0001. (I-N) Confocal time-lapse images of actin dynamics (I, K, M) and color-coded time projection of cell outlines from several time points (J, L, N) from FBCs expressing Lsp2-Gal4+UAS-LifeAct-GFP+control (I, J), +UAS-Cdc42-N17 (K, L) or +UAS-Rac1-N17 (M, N). Red arrowheads point at actin waves and blue doted lines outline FBCs. See also Movie 17. (O) Quantification of number of actin waves seen during 20min-long movies from (I, K, M) in FBCs expressing Lsp2-Gal4+UAS-LifeAct-GFP+control (n:34cells) or +UAS-Cdc42-N17 (n:26cells) or +UAS-Rac1-N17 (n:36cells). Kruskal-Wallis test with multiple comparisons, ***p=0.0001 and **** p<0.0001. (P, Q). Quantification of mean aspect ratio (P) and mean aspect ratio change over time (Q) from (I, K, M) of FBCs expressing Lsp2-Gal4+UAS-LifeAct-GFP+control or +UAS-Cdc42-N17. Control (n:18cells), Cdc42-N17 (n:14cells) and Rac1-N17 (n:15cells). Kruskal-Wallis test with multiple comparisons, * p:0.0141 and **** p<0.0001 (P). Ordinary one-way ANOVA test with multiple comparisons, **** p<0.0001 (Q). (R) Diagram of the proposed model for FBC swimming migration. Scale bars, 100 µm (A-D), 20 µm (F-H and J-L).

Cdc42 and Rac1 are known to regulate actin polymerisation during mesenchymal migration (Ridley, 2001). Hence, we tested if Cdc42 and Rac1 might regulate actin wave formation in FBCs. Indeed, Cdc42-N17 expression decreased the number of actin waves in FBCs dramatically compared to the control (Figure 6I, K, O and Movie 17A and B). Moreover, this loss of actin waves was associated with cell rounding alongside a strong reduction in cell shape deformations (Figure 6J, L, P and Q). Rac1-N17 expression also resulted in a strong, significant reduction in the number actin waves (Figure 6I, M, O and Movie 17A and C) and cell rounding as well as reduced cell shape changes (Figure 6J, N, P and Q) although the effects were less severe than upon Cdc42-N17 expression. Thus, Cdc42 and Rac1 are both major regulators of actin wave formation and cell deformations during FBC migration. Interestingly, we find that cell deformations (measured as aspect ratio change) directly correlated with migration speed in all the genetic manipulations affecting FBC migration that we tested (Suppl. Figure 3, Pearson test, *R*=0.7315). This further confirms that cell deformations are a major contributor to FBC migration.

In mesenchymal migration, Cdc42 and Rac1 are known to regulate Arp2/3-driven branched actin formation via Wasp and the Scar/Wave complex (Ridley, 2001, Ridley, 2015). Hence, we tested if these effectors of Cdc42 and Rac1 might also participate in FBC migration by inducing branched actin formation by expressing RNAi lines for Wasp, Arp2 and Arp3 specifically in FBCs. RNAi against Wasp as well as RNAi against Arp2 or Arp3 led to a significant decrease in FBC migration speed (Suppl. Figure 4A-C and H, and D-G and I, respectively). This suggests that Wasp and the Arp2/3 complex are involved in FBC migration and might be the effectors of Cdc42.

Altogether, here we use Drosophila FBCs as the first and currently only *in vivo* model of swimming migration to study the molecular mechanism underlying this unusual amoeboid migration mode. Our data show that, in the absence of a wound, FBCs patrol the pupal body by active swimming migration in hemolymph. For this, FBCs periodically generate dynamic actin waves which depends jointly on RhoA, Rac1 and Cdc42 and their effector Dia. The peristaltic movement of these actin waves leads to myosin-dependent cell surface compressions at the cell rear. As a result, the cells then elongate towards the opposite direction to drive cell locomotion.

## Discussion

Swimming migration of non-flagellated cells used to be considered a theoretically possible mode of cell migration until recent *in vitro* studies demonstrated that several cell types can indeed swim in solution (Barry and Bretscher, 2010, Garcia-Seyda et al., 2021, Aoun et al., 2020, O’Neill et al., 2018). These studies have made important progress in this field, by showing that membrane paddling coupled to a retrograde actin flow drives force transmission during swimming migration of macrophages and T cells (O’Neill et al., 2018, Aoun et al., 2020). However, the mechanism underlying swimming cell migration, in particular with regards to force generation, remained ill-defined. Moreover, swimming migration had not been studied *in vivo,* and hence the mechanism driving swimming migration under physiological conditions was completely unknown. Whether any cells even use swimming migration in *in vivo* environments was also unknown, until we recently described the swimming migration of pupal FBCs through hemolymph towards wounds (Franz et al., 2018). Here we show that, even in the absence of wounds, pupal FBCs actively move around the pupal body at high speed using swimming migration. However, instead of displaying directed migration, in this case they use random walk. Having established FBCs as the first *in vivo* model of swimming migration, we characterised the mechanism of force generation during swimming migration of FBCs during random walk in detail. For this, we developed an automatic cell tracking assay that allowed us to assess migration speed at large scale and in high-throughput and combined it with *in vivo* live imaging at high spatio-temporal resolution.

We find that FBCs have a unique cell morphology and migration characteristics compared to other motile cells described before. FBCs are very large, spherical cells filled with lipid droplets that, unlike most migrating cells, migrate without forming any protrusions at the cell front such as blebs, lamellipodia or filopodia. Instead, they maintain a round shape in the front throughout their migration. This is somewhat similar to embryonic progenitor cells in zebrafish that use amoeboid stable-bleb cell migration (Ruprecht et al., 2015), although these cells maintain a stable pear-like shape throughout their migration while FBCs undergo dynamic cell deformations.

*In vivo* swimming migration of FBCs is driven by periodic actin waves traveling to the cell rear. In contrast, *in vitro* swimming migration of leukocytes relies on a rearward cortical actin flow that is continuous (Aoun et al., 2020, O’Neill et al., 2018). In RAW macrophages, this rearward actin flow gets induced experimentally through local optogenetic RhoA activation which increases cortical actomyosin contractility (O’Neill et al., 2018). Similarly, amoeboid cell migration of several cell types is driven by continuous actin flows. These flows get induced through strong cell confinement which increases myosin II activity (Ruprecht et al., 2015, Liu et al., 2015, Logue et al., 2015). In contrast, FBCs *in vivo* do not appear to be under any such high confinement. Even in the more crowded areas such as the dorsal head and thorax we never observed any flattened cell surface areas indicative of the cell being compressed during FBC migration (Figure 1F and F’). Moreover, our previous study showed that FBCs also migrate with similar high speed and periodic actin waves in the much less crowded ventral part of the pupal thorax. This area is a big open space surrounded by epidermis which is almost entirely filled with hemolymph and where the small number of swimming FBCs are rarely in contact with a solid substrate ( (Franz et al., 2018) and Figure 1F’’). The lack of strong confinement in the pupa could explain why migratory FBCs lack a continuous actin flow, but rather have a dynamic actin network that periodically generates retrograde cortical actin waves. This also highlights that the environment is a key factor in dictating migration modes and the importance of studying cell migration under physiological settings.

*In vitro* swimming of leukocytes is driven by a constant retrograde actin flow which is regulated mainly by RhoA (Aoun et al., 2020, O’Neill et al., 2018). In contrast, we find here that *in vivo* swimming migration of FBCs, requires all three RhoGTPases, RhoA, Rac1 and Cdc42, to generate repetitive contractile actin waves. Actin wave formation is dependent on Dia-driven actin polymerization and, we suspect that Arp2/3-driven actin branching also contributes (Figure 6R). In addition, myosin II-driven actomyosin contraction might also contribute to actin wave formation possibly by pulling on the actin mesh to concentrate it. Indeed, when we blocked myosin II, we observed a decrease in the number of actin waves with the remaining waves appearing weaker (Figure 4P, Movie 12). Moreover, FBC migration seems to rely on a highly dynamic actin network. Indeed, the expression of Utrophin-GFP and constitutively active Dia led to a denser and more persistent actin network covering most of the cortex which resulted in reduced cell deformations and migration speed (Suppl. Figure 2A, D-F and H and Figure 5A, D, E, J, L and R). In addition, actin-depolymerizing proteins such as Cofilin might play a role in dissolving actin waves during FBC migration, since Cofilin is crucial for stable bleb amoeboid migration to maintain cortical actin flow (Ullo and Logue, 2021). How exactly FBCs regulate where and when actin waves form and to what extend actin polymerisation, actin branching, actin depolymerisation and F-actin turnover play a role, will be an interesting area of future studies.

Once an actin wave has been generated, there is a myosin II-dependent contraction of the actin meshwork at the cell rear regulated through RhoA and Rok activation. This, in turn, results in a belt-like constriction applying compressive forces at the rear which causes the cell to elongate towards the front resulting in cell translocation (Figure 6R).

Overall, we find that FBC migration is orchestrated by all three small RhoGTPases and several of their effectors. This possibly involves a complex crosstalk between the RhoGTPases to regulate actin wave formation and contraction, which will need to be deciphered in future work.

Interestingly, the finding that FBCs use repetitive actin waves rather than a stable cortical flow could explain the apparent stop-and-go or run-and-tumble mode of migration of FBCs (Suppl. Figure 1D). This might be caused by alternating between active migration phases when strong actin waves generate cell deformations to propel the cell body forward and more passive phases during which no, or weaker actin waves are formed. The use of repetitive actin waves might also provide more flexibility in changing direction to allow manoeuvring in the complex 3D environments found *in vivo*. This raises the question of how changes in direction get regulated. This might either be via a random repolarisation after each wave cycle, and/or might depend on some external signals, e.g. signals released by a wound, or by FBC-FBC contacts potentially inducing a repellent behaviour (“contact inhibition of locomotion”). Indeed, in the more crowded dorsal thorax we occasionally see FBCs transiently contacting neighbouring FBCs (Figure 1F’) which might affect the migration. However, we have not found any clear evidence of contact inhibition of locomotion playing a role in FBC spreading, such as a repulsive behaviour or the induction of actin waves upon FBC-FBC contact. Moreover, we do not believe that the transient FBC-FBC contacts we observe are generally required for FBC swimming migration, since FBCs swim with a similar speed in the much less crowded ventral thorax, where many FBCs migrate for long periods without forming any cell-cell contacts while only being surrounded by hemolymph ( (Franz et al., 2018) and Figure 1F’’). Together, this suggests, that the efficient spreading of FBCs in the pupal body is driven by a sporadic, cell-intrinsic mechanism of random walk rather than contact inhibition of locomotion.

An intriguing question that arises from swimming migration, is how the forces that are generated inside the cell, get transmitted to the aqueous environment to propel the cell body forward. In mesenchymal migration this force transmission is accomplished by cell-ECM adhesion to allow traction. However, swimming cell migration does not involve cell-ECM adhesion. Two models of force transmission in swimming migration have been proposed: nonreciprocal cell shape changes or cell membrane treadmilling (Paluch et al., 2016). *In vitro* studies of swimming migration of lymphocytes and macrophages showed that a rearward flow of plasma membrane proteins (“membrane paddling”) coupled to the cortical actin flow is critical for swimming migration (Aoun et al., 2020, O’Neill et al., 2018). In contrast, cell deformations appeared to play only a minor role. The mechanism of force transmission in FBCs migration is still completely unknown and will be the focus of future studies.

Here we describe for the first time an *in vivo* swimming cell migration mode that allows the fast travel of FBCs within the aqueous environment of the pupa over long distances. This raises the intriguing question of whether this swimming migration mode might enable other cells, not just in invertebrates but potentially also in vertebrates, to perform fast efficient spreading when encountering certain liquid filled environments such as confining conjunctive tissues or liquid-rich infected areas such as oedemas in chronic wounds. This might be particularly useful for cells that need to travel across different body parts such as immune cells and cancer cells during cancer metastasis.

## Materials and Methods

### Fly stocks and maintenance

Drosophila melanogaster stocks and crosses were maintained and performed on cornmeal molasses food at 25°C. The following lines were used in this paper: w^67^ as a control, Lsp2-Gal4 (BDSC: 6357), UAS-NLS-mCherry (BSDC: 38424), UAS-LifeAct-GFP (Zanet et al., 2012), UAS-LifeAct-Scarlet (Serna-Morales et al., 2023), UAS-GMA-GFP(Dutta et al., 2002), UAS-Utrophin-GFP(Rauzi et al., 2010), UAS-Rho1-GFP (BSDC: 9393), Ubi>Anillin-RBD-GFP(Munjal et al., 2015), sqh^AX3^;sqh>Sqh-GFP (BSDC: 57144), sqh>Rok^K116A^-Venus (Bulgakova et al., 2013), Lpp-GFP (VDRC: FTRG-318255), Ubi>CaaX-RFP (DGRC: 109826), UAS-Zipper-DN-GFP (Monier et al., 2010), UAS-Rho1-N19 (BSDC: 7327), UAS-Rho1-RNAi (gift from Brian Stramer), UAS-Rok-RNAi 1 (BSDC: 28797) and 2 (VDRC: KK-104675), UAS-Sqh-RNAi (VDRC: KK-109493), UAS-Sqh-AA (BSDC: 64114), UAS-Dia RNAi 1 (BSDC: 28541) and 2 (VDRC: KK-103914), UAS-DiaΔDAD-GFP (BSDC: 56752), UAS-CDC42-N17 (BSDC: 6288), UAS-CDC42-RNAi (BSDC: 29004), UAS-Rac1-N17 (BSDC: 6292), UAS-Rac1-RNAi (BSDC: 28985), UAS-WASp-RNAi 1 (BSDC: 25955) and 2 (VDRC: KK-109220), UAS-Arp2-RNAi 1 (VDRC: KK-101999) and 2 (VDRC: GD-29944) and UAS-Arp3-RNAi (VDRC: GD-35260).

### Microscopy movies and images

Pupae were aged to the developmental age of 16h after pupal formation at 25°C by marking white pre-pupae (0h). Pupae were dissected at 16h APF by removing the pupal case (Weavers et al., 2018) and placed on a coverslip on their dorsal side or on their head.

Movies of whole pupae used to track FBCs migration were collected on a Zeiss Celldiscovery 7 widefield microscope with a 5X lens at 0.5 magnification for 3h with a time interval of 1min30sec (Figure 1A-B and Movie 1) or 5min (Figures 3-6, Suppl. Figures 2 and 4, and Movies 6, 9, 13 and 16). Movies to follow actin waves and FBC deformations were acquired with a Nikon SoRa spinning disk confocal microscope using a 40X silicon objective for 20min with a time interval of 30sec using a magnification of 1. ‘Higher resolution’ movies (Figures 2F-F’ and 3E, Suppl. Figure 2F-H and Movies 5, 7 and 11) were acquired with the Nikon SoRa spinning disk confocal microscope with a 40X silicon objective using a magnification of 2.8. Images in figure 3E, 4F and movies 7A and 10A were obtained with a Nikon AX-R NSPARC. We deconvolved movies 5, 7A and 10B with the Nikon NIS-Elements imaging software using a Richardson-Lucy deconvolution method with 9-13 iterations. Movies 2 and 10 were acquired with a Zeiss LSM 980 confocal microscope using a 63X oil objective.

Methylene blue-stained, semi-thick serial transverse sections of 1μm thickness (Figure 1F), were prepared as described in (Franz et al., 2018) and a Leica MICA with a 20x objective was used to collect images.

FBC tracking in movies of pupae was automatically generated using IMARIS software. Only 1h30-3h long uninterrupted tracks (∼100 per pupa) were analysed and are shown in images as whole tracks and in movies as dragon tail tracks. Images and movies of confocal movies were generated with FIJI ImageJ to create z-projections, z-sections and orthogonal view images. We used the same brightness and contrast adjustment for control and experimental conditions. Movies and images were organised and annotated with VSDC Video Editor and Abode Illustrator.

### Analysis

#### Migration tracking

FBCs nuclear tracking was generated blindly and automatically using IMARIS software in the dorsal head, thorax and abdomen of pupae. The tracking was obtained in 2D to avoid artefacts created in the z-axis due to poorer z resolution. Only tracks with a duration of 1h30min-3h were selected and analysed. We then examined visually if the tracks were correct and were corrected or deleted incorrect tracks. Average speed or straightness per pupa were obtained by first obtaining the mean speed or straightness of each individual track in a certain pupa (mean over 1h30min-3h depending on track length) and then calculating the mean speed or straightness from these mean values. Tracks were color-coded based on their average speed or duration or speed over time with IMARIS.

#### Actin wave number and FBC deformation analysis

Confocal movies were analysed in FIJI ImageJ. We quantified the number of actin waves and cell deformation only in FBCs which were fully in the field of view throughout the 20min-long movies. Visual assessment of actin wave numbers was done blindly. Actin waves were defined as a local cortical increase in Lifeact-GFP intensity followed by a decrease. For the aspect ratio, we drew the outline around a particular FBC every minute and measured the aspect ratio. Next, we calculated the mean aspect ratio and the mean change in aspect ratio, which is the absolute value of the difference in aspect ratio between one minute and the next. All these quantifications were done blindly.

Concave and convex index images were generated in FIJI Image J by manually segmenting FBCs and using the FIJI Image J plugging Adapt (Barry et al., 2015).

### Quantification of contact sites of FBCs with other tissues

We quantified the length of the FBC border in contact with the hemolymph, other FBCs, epidermis, hemocytes and other pupal tissues by using FIJI. We analysed only FBCs which were fully contained within the 15-20 serial sections analysed. The percentage of FBC surface contacting other tissues was calculated for each section by dividing the length of the FBC border in contact with another tissue by the total length of the FBC outline and the mean was then calculated from all sections for each cell.

### Statistics

Statistics were performed using GraphPad Prism 10. Gaussian distribution was assessed to test datasets. We used parametric or non-parametric tests depending on the Gaussian or non-Gaussian characteristics of the data distribution. P<0.05 was set as the significance threshold. For the box and whisker plots, the median is plotted as line inside the box. The box extends from the 25^th^ to the 75^th^ percentile, and the whole dataset is shown by the whiskers and dots. In scatter dot plots, the line in the middle indicates the median and the error bars show the extents of the whole dataset.

## Acknowledgements

We would like to thank members of the Amoyel, Fernandes, Mao, Pichaud and Stramer labs for helpful discussion. Thanks to Marc Amoyel (UCL), Marcus Bischoff (University of St Andrews), Vilaiwan Fernandes (UCL), Yanlan Mao (UCL), Isabel Palacios (Queen Mary University), Brian Stramer (King’s College London), the Vienna Drosophila Resource Center (Vienna, Austria) and the Bloomington Drosophila Stock Center (Indiana, USA) for *Drosophila* stocks. We are also thankful to Barbara Conradt, Roberto Mayor and Eric Theveneau for critical comments on the manuscript. Finally, a big thanks goes to the team of the UCL Biosciences Imaging Facility for their help with imaging and image analysis. This work was funded by the Wellcome Trust and Royal Society Sir Henry Dale fellowship granted to A.F. (215431/Z/19/Z).

## Author contributions

C.A. designed and performed the experiments with the help of B.H.L., C.A. and A.F. designed the study and C.A. and A.F. wrote the manuscript.

