## Supplementary data + Movie Legends for "Cell deformations generated by dynamic cortical actin waves drive *in vivo* swimming migration"

### Supplementary Figure 1

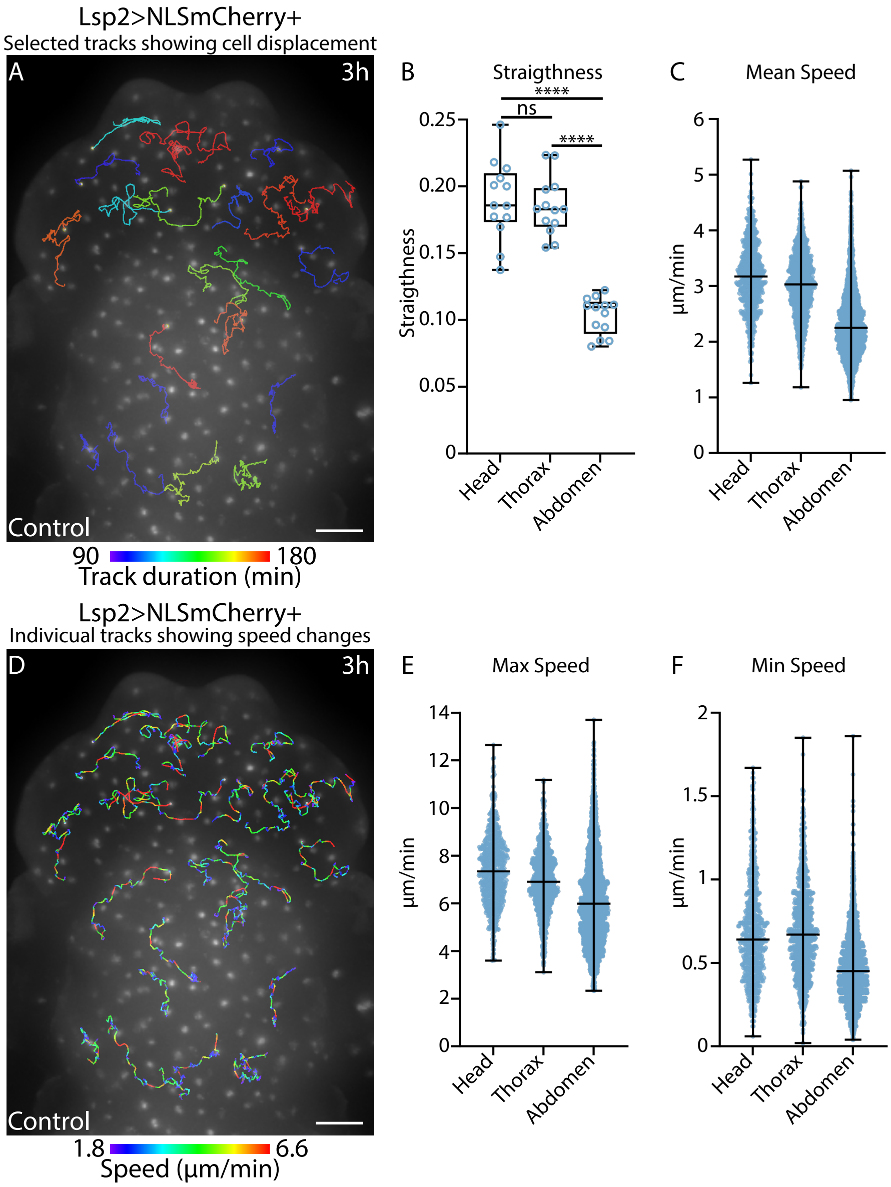

Supplementary Figure 2

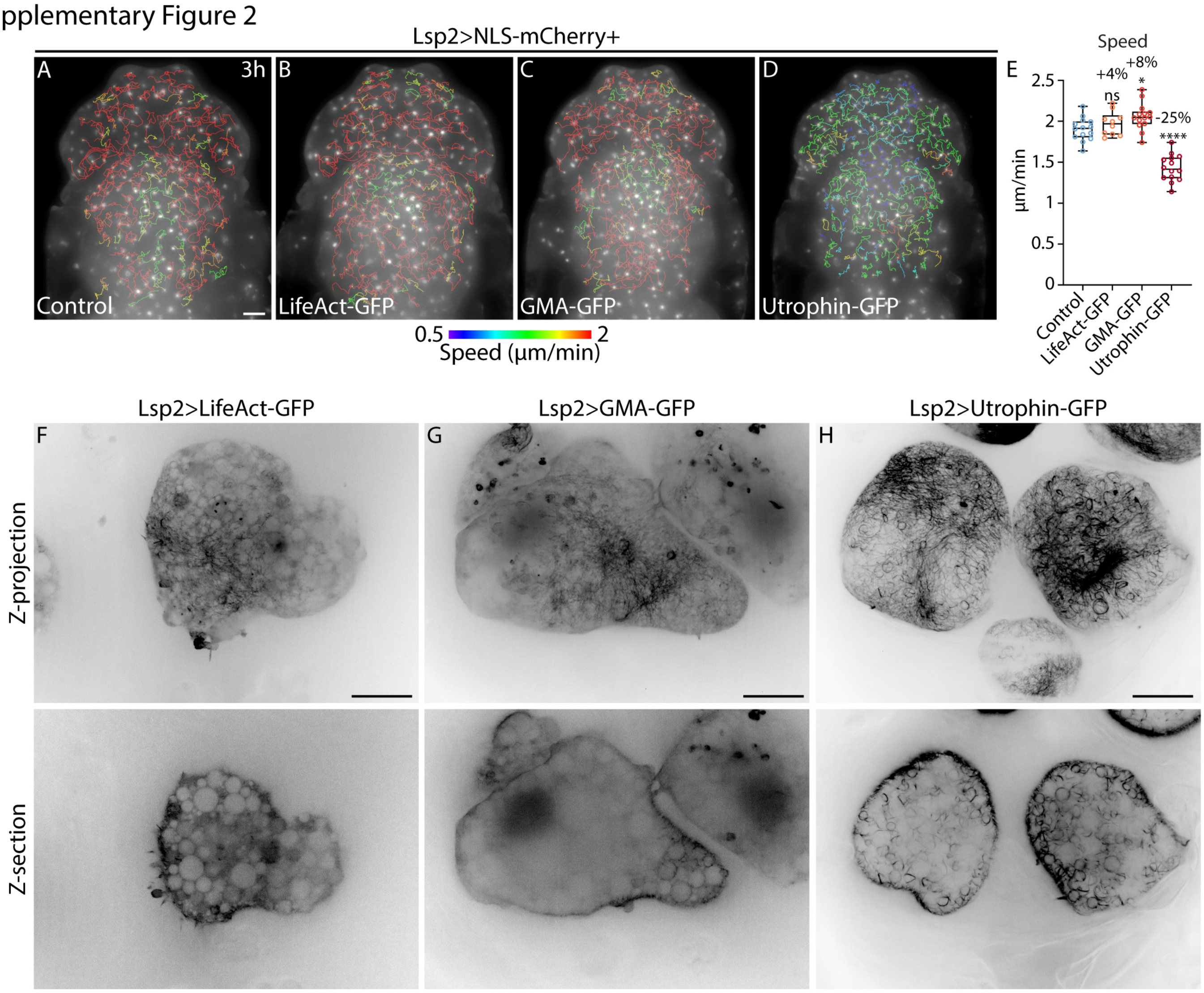

Supplementary Figure 3

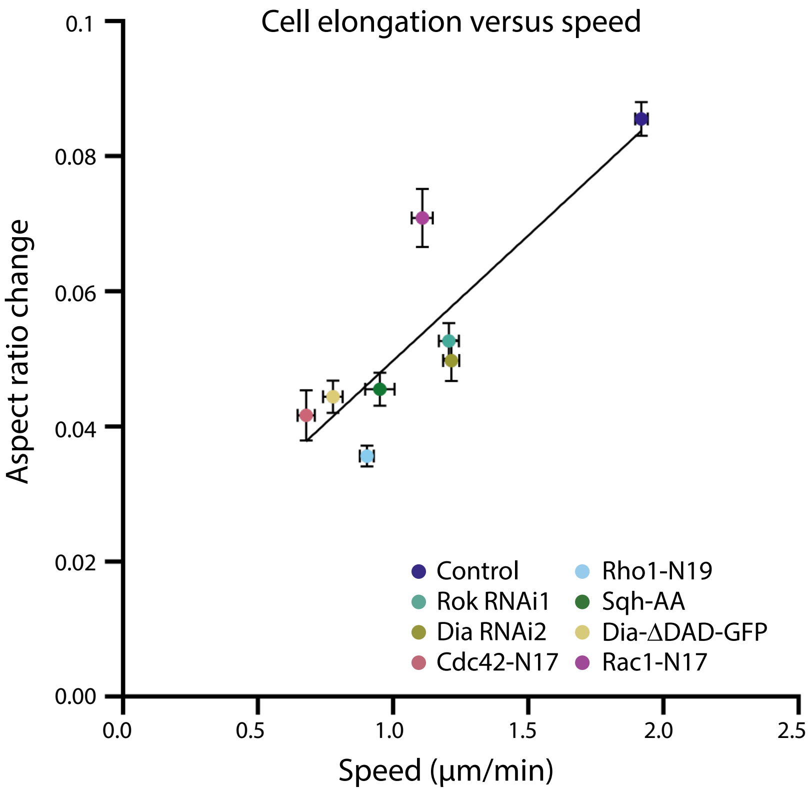

Supplementary Figure 4

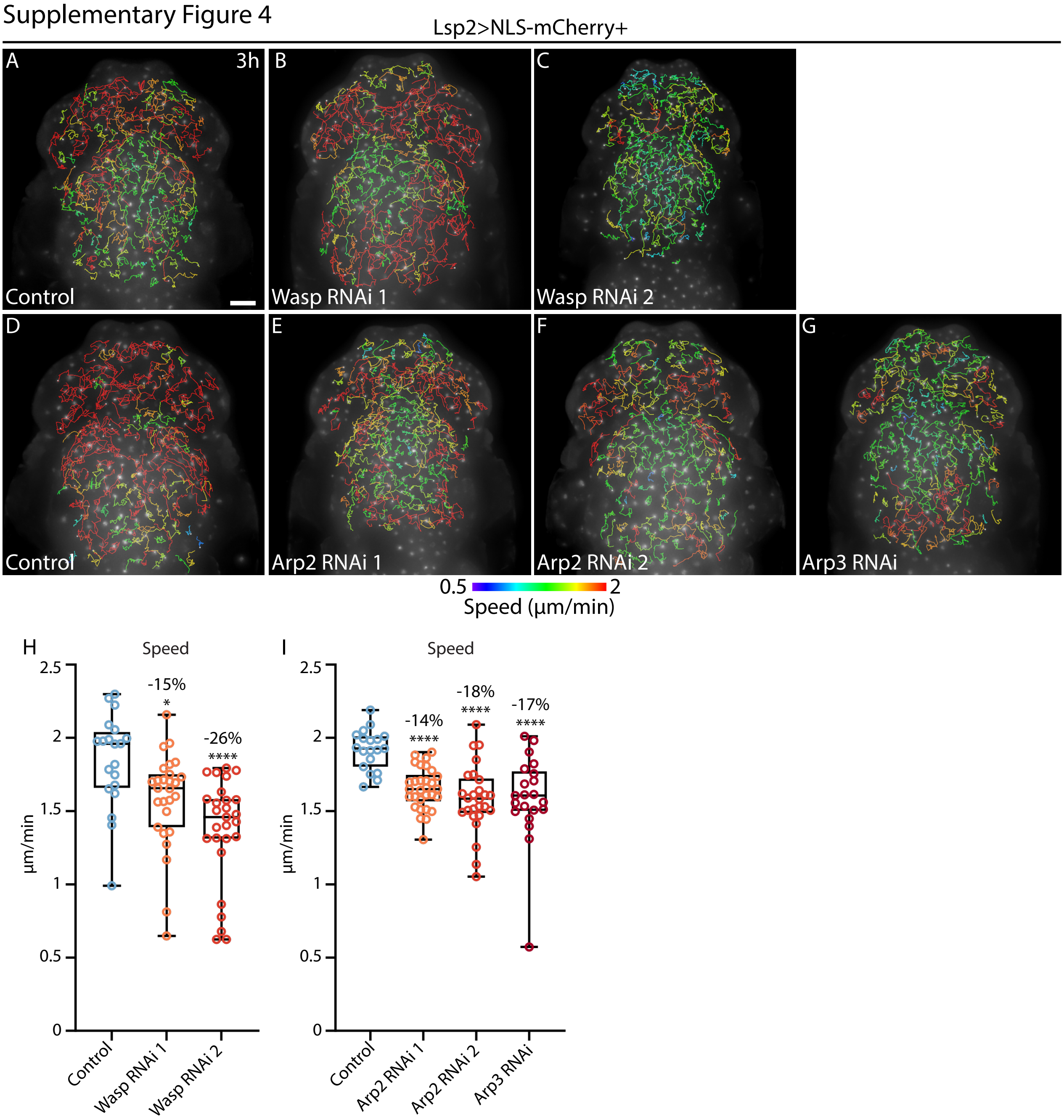

| Figure | Genotype |
| --- | --- |
| 1A | ;; Lsp2-Gal4+UAS-NLS-mCherry/+ |
| 1B | ; UAS-Zipper-DN-GFP/+ ; Lsp2-Gal4+UAS-NLS-mCherry/+ |
| 1C | ;; Lpp-GFP+Ubi>CAAX-RFP/Lsp2-Gal4+UAS-NLS-mCherry |
| 2 | ;; Lsp2-Gal4+UAS-LifeAct-GFP+UAS-NLS-mCherry/+ |
| 3A | ;; Lsp2-Gal4+UAS-NLS-mCherry/+ |
| 3B | UAS-Rho1-N19/+ or Y ;; Lsp2-Gal4+UAS-NLS-mCherry/+ |
| 3C | ;; Lsp2-Gal4+UAS-NLS-mCherry/UAS-Rho1 RNAi |
| 3E | ; UAS-Rho1-GFP/+ ; Lsp2-Gal4+UAS-NLS-mCherry/+ |
| 3F | ; Ubi>Ani-RBD-GFP/Ubi>Ani-RBD-GFP ; |
| 3G | ;; Lsp2-Gal4+UAS-LifeAct-GFP+UAS-NLS-mCherry/+ |
| 3I | UAS-Rho1-N19/+ ;; Lsp2-Gal4+UAS-LifeAct-GFP+UAS-NLS-mCherry/+ |
| 4A | ;; Lsp2-Gal4+UAS-NLS-mCherry/+ |
| 4B | ;; Lsp2-Gal4+UAS-NLS-mCherry/UAS-Rok RNAi 1 |
| 4C | ; UAS-Rok RNAi 2/+ ; Lsp2-Gal4+UAS-NLS-mCherry/+ |
| 4D | ;; Lsp2-Gal4+UAS-NLS-mCherry/UAS-Sqh-AA |
| 4E | ; UAS-Sqh RNAi ; Lsp2-Gal4+UAS-NLS-mCherry/+ |
| 4F | ; UAS-Zipper-DN-GFP/+ ; Lsp2-Gal4+UAS-NLS-mCherry/+ |
| 4H | ;; Sqh>Rok$^{K116A}$-Venus/ Sqh>Rok$^{K116A}$-Venus |
| 4I | Sqh$^{Ax3}$;;Sqh>Sqh-GFP/Lsp2-Gal4+UAS-NLS-mCherry |
| 4J | ;; Lsp2-Gal4+UAS-LifeAct-GFP+UAS-NLS-mCherry/+ |
| 4L | ;; Lsp2-Gal4+UAS-LifeAct-GFP+UAS-NLS-mCherry/UAS-Rok RNAi 1 |
| 4N | ;; Lsp2-Gal4+UAS-LifeAct-GFP+UAS-NLS-mCherry/UAS-Sqh-AA |
| 5A | ;; Lsp2-Gal4+UAS-NLS-mCherry/+ |
| 5B | ;; Lsp2-Gal4+UAS-NLS-mCherry/UAS-Dia RNAi 1 |
| 5C | ; UAS-Dia RNAi 2/+ ; Lsp2-Gal4+UAS-NLS-mCherry/+ |
| 5D | ;; Lsp2-Gal4+UAS-NLS-mCherry/UAS-Dia$\Delta$DAD-GFP |
| 5F | ;; Lsp2-Gal4+UAS-LifeAct-GFP+UAS-NLS-mCherry/+ |
| 5H | ; UAS-Dia RNAi 2/+ ; Lsp2-Gal4+UAS-LifeAct-GFP+UAS-NLS-mCherry/+ |
| 5J | ;; Lsp2-Gal4+UAS-LifeAct-Scarlet /+ |
| 5L | ;; Lsp2-Gal4+UAS-LifeAct-Scarlet/UAS-Dia$\Delta$DAD-GFP |
| 6A | ;; Lsp2-Gal4+UAS-NLS-mCherry/+ |
| 6B | ; UAS-Cdc42-N17/+ ; Lsp2-Gal4+UAS-NLS-mCherry/+ |
| 6C | UAS-Cdc42 RNAi/+ ;; Lsp2-Gal4+UAS-NLS-mCherry/+ |
| 6E | ;; Lsp2-Gal4+UAS-NLS-mCherry/+ |
| 6F | ;; Lsp2-Gal4+UAS-NLS-mCherry/UAS-Rac1-N17 |
| 6G | ;; Lsp2-Gal4+UAS-NLS-mCherry/UAS-Rac1 RNAi |
| 6I | ;; Lsp2-Gal4+UAS-LifeAct-GFP+UAS-NLS-mCherry/+ |
| 6K | ; UAS-CDC42-N17/+ ; Lsp2-Gal4+UAS-LifeAct-GFP+UAS-NLS-mCherry/+ |
| 6M | ;; Lsp2-Gal4+UAS-LifeAct-GFP+UAS-NLS-mCherry/UAS-Rac1-N17 |
| S1 | ;; Lsp2-Gal4+UAS-NLS-mCherry/+ |
| S2A | ;; Lsp2-Gal4+UAS-NLS-mCherry/+ |

| | |
|---|---|
| S2B and F | *;; Lsp2-Gal4+UAS-NLS-mCherry/UAS-LifeAct-GFP* |
| S2C and G | *;; Lsp2-Gal4+UAS-NLS-mCherry/UAS-GMA-GFP* |
| S2D and H | *;; Lsp2-Gal4+UAS-NLS-mCherry/UAS-Utrophin-GFP* |
| S4A | *;; Lsp2-Gal4+UAS-NLS-mCherry/+* |
| S4B | *;; Lsp2-Gal4+UAS-NLS-mCherry/UAS-Wasp RNAi 1* |
| S4C | *; UAS-Wasp RNAi 2 ; Lsp2-Gal4+UAS-NLS-mCherry/+* |
| S4D | *;; Lsp2-Gal4+UAS-NLS-mCherry/+* |
| S4E | *; UAS-Arp2 RNAi 1; Lsp2-Gal4+UAS-NLS-mCherry/+* |
| S4F | *;; Lsp2-Gal4+UAS-NLS-mCherry/ UAS-Arp2 RNAi 2* |
| S4G | *;; Lsp2-Gal4+UAS-NLS-mCherry/UAS-Arp3 RNAi* |
| Movie 11 | $Sqh^{Ax3}$*;;Sqh>Sqh-GFP/Lsp2-Gal4+UAS-LifeAct-Scarlet* |

**Supplementary Figure 1 – Fat body cell speed varies during migration**

(A) Widefield timelapse images of the dorsal head and thorax of *Drosophila* pupae expressing Lsp2-Gal4+UAS-NLS-mCherry+control from Figure 1A and Movie 1A showing only a selection of the 1h30-3h tracks (shown color-coded according to track duration) to visualise the total distance that individual FBCs travel inside the pupa.

(B) Quantification of mean straightness of the tracks of migrating FBC expressing Lsp2-Gal4+UAS-NLS-mCherry+control in the head, thorax and abdomen (n:13pupae). Showing mean straightness for each pupa calculated from the mean straightness of each of its 1h30-3h-long tracks. Ordinary one-way ANOVA test with multiple comparisons, ns p:0.9302 and **** p<0.0001.

(C) Quantification of mean FBC speed in the head, thorax and abdomen of *Drosophila* pupae expressing Lsp2-Gal4+UAS-NLS-mCherry+control. Showing mean speed of all tracks of FBCS in the head (n:659cells), thorax (n:989cells) and abdomen (n:2366cells) from 13 pupae.

(D) Widefield timelapse images of the dorsal head and thorax of *Drosophila* pupae expressing Lsp2-Gal4+UAS-NLS-mCherry+control from (A) showing the same selection of 1h30-3hs tracks but color-coded by speed over time.

(E-F) Quantification of maximum speed (E) and minimum speed (F) from each FBC track in the head, thorax and abdomen of *Drosophila* pupae expressing Lsp2-Gal4+UAS-NLS-mCherry+control. Showing data from all tracks of FBCS in the head (n:659cells), thorax (n:989cells) and abdomen (n:2366cells) from 13 pupae.

Scale bars, 100 μm.

**Supplementary Figure 2– LifeAct labels actin meshwork without affecting fat body cell migration.**

(A-D) Widefield timelapse images of the dorsal head and thorax of *Drosophila* pupae expressing Lsp2-Gal4+UAS-NLS-mCherry+control (A), +UAS-LifeAct-GFP (B), +UAS-GMA-GFP (C) or +UAS-Utrophin-GFP (D). 1h30-3h-long migration tracks are shown color-coded according to their mean speed.

(E) Quantification of mean FBC speed from (A-D). Control (n:14pupae), LifeAct-GFP (n:10pupae), GMA-GFP (n:14pupae) and Utrophin-GFP (n:14pupae). Showing mean speed for each pupa calculated from the mean speed of each of its 1h30-3h-long tracks. One-way ANOVA test with multiple comparisons, ns p=0.591, *p=0.0293 and ****p<0.0001.

(F-H) Confocal time-lapse images of actin in FBCs expressing Lsp2-Gal4+UAS-LifeAct-GFP (F), +UAS-GMA-GFP (G) or +UAS-Utrophin-GFP (H). Z-projection in the top and Z-section in the bottom.

Scale bars, 100 μm (A-D), 20 μm (F-H).

**Supplementary Figure 3 – Cell deformations are a major contributor to fat body cell migration.**

Fat body cell aspect ratio change as a function of fat body cell speed. Mean FBC speed values of pupae expressing Lsp2-Gal4+UAS-NLS-mCherry+control (pooled from Figures 3D, 4G, 5E and 6D and H), +UAS-Rho1-N19 (Figure 3D), +UAS-Rok RNAi1 (Figure 4G), +UAS-Sqh-AA (Figure 4G), +UAS-Dia RNAi 2 (Figure 5E), +UAS-Dia-ΔDAD-GFP (Figure 5E), +UAS-Cdc42-N17 (Figure 6D) and +UAS-Rac1-N17 (Figure 6H). Aspect ratio change values of pupae expressing Lsp2-Gal4+UAS-LifeAct-GFP+control or Lsp2-Gal4+UAS-LifeAct-Scarlet+control (pooled from Figures 3M, 4R, 5P and 6Q), +UAS-Rho1-N19 (Figure 3M),+UAS-Rok RNAi1 (Figure 4R), +UAS-Sqh-AA (Figure 4R), +UAS-Dia RNAi 2 (Figure 5P), +UAS-Dia-ΔDAD-GFP (Figure 5R), +UAS-Cdc42-N17 (Figure 6Q) and +UAS-Rac1-N17 (Figure 6Q). Pearson test, $R$=0.7315.

**Supplementary Figure 4 – Wasp and the Arp2/3 complex are involved in fat body cell migration.**

(A-G) Widefield timelapse images of the dorsal head and thorax of *Drosophila* pupae expressing Lsp2-Gal4+UAS-NLS-mCherry+control (A, D), +UAS-Wasp RNAi1 (B), +UAS-Wasp RNAi2 (C), +UAS-Arp2 RNAi1 (E), +UAS-Arp2 RNAi2 (F) and +UAS-Arp3 RNAi (G). 1h30-3h-long migration tracks are shown color-coded according to their mean speed.

(H, I) Quantification of mean FBC speed from (A-G). Control (n:20pupae), Wasp RNAi1 (n:27pupae) and Wasp RNAi2 (n:27pupae) (H) and control (n:19pupae), Arp2 RNAi1 (n:30pupae), Arp2 RNAi2 (n:26pupae) and Arp3 RNAi (n:21pupae) (I). Showing mean speed for each pupa calculated from the mean speed of each of its 1h30-3h-long tracks. Kruskal-Wallis test with multiple comparisons, *p = 0.0172 and ****p<0.0001.

Scale bar, 100 μm.

**Movie 1 - Fat body cells migrate in the whole pupa – Related to Figure 1A-B.**

Widefield movies of the dorsal view of *Drosophila* pupae expressing Lsp2-Gal4+UAS-NLS-mCherry+control (A) or +UAS-Zipper-DN-GFP (B). Only continuous 1h30-3h long migration tracks with a dragon-tail are shown color-coded according to their mean speed. Elapsed time is in top left corner in minutes:seconds. Scale bar, 100 μm.

**Movie 2 - Fat body cells swim inside the hemolymph – Related to Figure 1E.**

Confocal movie of two FBCs swimming in the hemolymph near the epidermis in a Lpp-GFP+Ubi>CaaX-RFP+Lsp2-Gal4+UAS-NLS-mCherry pupa (hemolymph in green; epidermis in magenta on the top; FBCs in faint magenta with bright magenta nuclei, note presence of a hemocyte containing a bright magenta object). Elapsed time is in top left corner in minutes:seconds. Scale bar, 20 μm.

**Movie 3 - Actin waves deforming fat body cell during swimming migration – Related to Figure 2A.**

Confocal movie of actin dynamics in migrating FBCs expressing Lsp2-Gal4+UAS-LifeAct-GFP. Red arrowheads point at consecutive actin waves in the rear of a migrating FBC. Elapsed time shown in top left corner in minutes:seconds. Scale bar, 20 µm.

**Movie 4 - Actin wave leads to local deformation of fat body cell rear – Related to Figure 2D.**

Confocal movie a Lsp2-Gal4+UAS-LifeAct-GFP expressing FBC. Red arrowheads show actin wave moving to the rear. Blue arrows point at FBC cell surface compressions. Elapsed time shown in top left corner in minutes:seconds. Scale bar, 20 µm.

**Movie 5 - High resolution of actin wave – Related to Figure 2F.**

'Higher resolution' confocal movie of a Lsp2-Gal4+UAS-LifeAct-GFP expressing FBC followed by a magnified version of the left top part of the FBC. Red arrowheads show actin wave movement. Elapsed time shown in top left corner in minutes:seconds. Scale bar, 20 µm.

**Movie 6 - Rho1 is crucial for fat body cell migration – Related to Figure 3A-C.**

Widefield movies of the dorsal head and thorax of *Drosophila* pupae expressing Lsp2-Gal4+UAS-NLS-mCherry+control (A), +UAS-Rho1-N19 (dominant negative form of Rho1) (B) or +UAS-Rho1 RNAi (C). Only continuous 1h30-3h long migration tracks with a dragon-tail are shown color-coded according to their mean speed. Elapsed time shown in top left corner in hours:minutes. Scale bar, 100 µm.

**Movie 7 - Rho1 accumulates in fat body cell rear during migration – Related to Figure 3E-F.**

Confocal movies of a swimming FBC expressing Lsp2-Gal4+UAS-Rho1-GFP (A) or Ani-RBD-GFP under the control of the ubiquitin p63E promoter (B). Red arrowhead points at the accumulation of Rho1-GFP or Ani-RBD-GFP as punctae in the rear of a migratory FBC. Yellow dotted line outlines a hemocyte. Left panel is the maximum projection of the total Z-stack and right panel is one Z-section. Elapsed time shown in top left corner in minutes:seconds. Scale bar, 20 µm.

**Movie 8 - Rho1 is crucial for actin waves and fat body cell deformations – Related to Figure 3G-J.**

Confocal movies of actin dynamics and cell deformation from FBCs expressing Lsp2-Gal4+UAS-LifeAct-GFP+control (A) or +UAS-Rho1-N19 (B). Red arrowhead points at actin wave. Note that white line in (A) is the shadow formed by a cuticle fold. Elapsed time shown in top left corner in minutes:seconds. Scale bar, 20 µm.

**Movie 9 - Actomyosin contractions are essential for fat body cell migration – Related Figure 4A-F.**

Widefield movies of the dorsal head and thorax of *Drosophila* pupae expressing Lsp2-Gal4+UAS-NLS-mCherry+control (A), +UAS-Rok RNAi1 (B), +UAS-Rok RNAi2 (C), +UAS-Sqh-AA (DN) (D), +UAS-Sqh RNAi (E) or +UAS-Zipper-DN-GFP (F). Only continuous 1h30-3h long migration tracks with a dragon-tail are shown color-coded according to their mean speed. Elapsed time shown in top left corner in hours:minutes. Scale bar, 100 µm.

**Movie 10 - Rok and myosin II are found at the rear of a migrating fat body cell – Related to Figure 4H-I.**

Confocal movie of migrating FBCs expressing $Rok^{K116A}$-Venus under the sqh promoter (A) and Sqh-GFP under the sqh promoter in a sqh mutant background (B). Red arrowhead points the accumulation of $Rok^{K116A}$-Venus or Sqh-GFP in the rear of a migrating FBC. For Rok-Venus, left panel is the maximum projection of the total Z-stack and right panel is the maximum projection of 4 deeper Z-sections. Elapsed time shown in top left corner in minutes:seconds. Scale bar, 20 µm.

**Movie 11 - Myosin II is closely associated with the actin in a wave.**

Higher resolution confocal movie across a Z stack of a FBC expressing Sqh-GFP under the sqh promoter in a sqh mutant background and UAS-LifeAct-Scarlet using the Lsp2-Gal4 driver. Magnified view in the top right corner. Scale bar, 20 µm, 5 µm (magnified view).

**Movie 12 - Rok and myosin II are needed for fat body cell deformations – Related to Figure 4J-O.**

Confocal movies of actin dynamics and cell deformation from FBCs expressing Lsp2-Gal4+UAS-LifeAct-GFP+control (A), +UAS-Rok RNAi1 (B) or +UAS-Sqh-AA (DN) (C). Red arrowhead points at actin wave. Elapsed time shown in top left corner in minutes:seconds. Scale bar, 20 µm.

**Movie 13 - Dia is involved in fat body cell migration – Related to Figure 5A-D.**

Widefield movies of the dorsal head and thorax of *Drosophila* pupae expressing Lsp2-Gal4+UAS-NLS-mCherry+control (A), +UAS-Dia RNAi1 (B), +UAS-Dia RNAi2 (C) or +UAS-Dia-ΔDAD-GFP (CA) (D). Only continuous 1h30-3h long migration tracks with a dragon-tail are shown color-coded according to their mean speed. Elapsed time shown in top left corner in hours:minutes. Scale bar, 100 µm.

**Movie 14 - Dia is involved in actin wave formation – Related to Figure 5F-I.**

Confocal movies of actin dynamics and cell deformation from FBCs expressing Lsp2-Gal4+UAS-LifeAct-GFP+control (A) or +UAS-Dia RNAi2 (B). Red arrowhead points at actin wave. Elapsed time shown in top left corner in minutes:seconds. Scale bar, 20 µm.

**Movie 15 - Dia overactivation increases cortical actin meshwork – Related to Figure 5J-M.**

Confocal movies of actin dynamics and cell deformation from FBCs expressing Lsp2-Gal4+UAS-LifeAct-Scarlet+control (A) or +UAS-Dia-ΔDAD-GFP (CA) (B). Red arrowhead points at actin waves (A) or highly dynamic actin swirls (B). Elapsed time shown in top left corner in minutes:seconds. Scale bar, 20 µm.

**Movie 16 - Cdc42 and Rac1 are crucial for fat body cell migration – Related to Figure 6A-C, E-G.**

Widefield movies of the dorsal head and thorax of *Drosophila* pupae expressing Lsp2-Gal4+UAS-NLS-mCherry+control (A and D), +UAS-Cdc42-N17 (DN) (B), +UAS-Cdc42 RNAi (C), +UAS-Rac1-N17 (DN) (E) or +UAS-Rac1-RNAi (F). Only continuous 1h30-3h long migration tracks with a dragon-tail are shown color-coded according to their mean speed. Elapsed time shown in top left corner in hours:minutes. Scale bar, 100 µm.

**Movie 17 - Cdc42 and Rac1 are involved in actin wave formation and fat body cell deformations – Related to Figure 6I-N.**

Confocal movies of actin dynamics and cell deformation from FBCs expressing Lsp2-Gal4+UAS-LifeAct-GFP+control (A), +UAS-Cdc42-N17 (DN) (B) or +UAS-Rac1-N17 (DN) (C). Red arrowhead points at actin wave. Elapsed time shown in top left corner in minutes:seconds. Scale bar, 20 µm.
